# 3D Electron Microscopy Reveals Diverse Chromosome Morphologies Across Dinoflagellate Species

**DOI:** 10.64898/2026.08.10.743404

**Authors:** Lucas Philipp, Eran Ittah, Dirk Schumann, Joachim de Fourestier, Natalie Reznikov, Stephanie C. Weber

**Affiliations:** Quantitative Life Sciences Program, McGill University, Montreal H3A 1E3, Canada; Department of Bioengineering, McGill University, Montreal H3A 0E9, Canada; Fibics Incorporated, Ottawa K2E 0B9, Canada; Department of Anatomy and Cell Biology, School of Biomedical Sciences, Faculty of Medicine and Health Sciences, Faculty of Dental Medicine and Oral Health Sciences, McGill University, Montreal H3A 0C7, Canada; Centre de Recherche en Biologie Structurale, McGill University, Montreal H3G 0B1, Canada; Departments of Biology and Physics, McGill University, Montreal H3A 1B1, Canada

**Keywords:** dinoflagellate, FIB-SEM, chromosome structure, species diversity, shape analysis, DNA toroid

## Abstract

Dinoflagellate chromosomes adopt a highly condensed and organized morphology, with periodic bands and arches observed by traditional Transmission Electron Microscopy (TEM). However, the limited two-dimensional field of view of TEM has prevented a precise characterization of the inherently three-dimensional organization of dinoflagellate chromosomes. Moreover, given the vast diversity among dinoflagellate species and the lack of a systematic comparison of their chromosomes, it remains unclear whether dinoflagellate chromosomes share common organizational features or instead exhibit significant cell- or species-specific differences. Here, we acquire three whole-nucleus 3D Focused Ion Beam Scanning Electron Microscopy (FIB-SEM) datasets at 4 nm voxel size for each of four dinoflagellate species: *Symbiodinium microadriaticum, Breviolum minutum, Fugacium kawagutii*, and *Crypthecodinium cohnii*. We compile these data with previously published image volumes from four additional species and present an analysis of the largest collection of dinoflagellate FIB-SEM images to date. Common features observed across all eight species include the absence of physical confinement or spatial clustering of chromosomes in the nucleus. In addition, by decomposing each chromosome into a weighted sum of orthogonal shapes using Spherical Harmonics Expansion, we find a principal component encapsulating 88% of the total shape variance that is common to all species. However, our analysis also reveals differences in chromosome morphology across species. First, while many chromosomes exhibit surface ridges with left-handed helical twist, the proportion of chromosomes with such ridges varies extensively across species. Second, while chromosomes in most species are discrete and well-separated, chromosomes in *F. kawagutii* are interconnected in a single contiguous network. Lastly, to our knowledge, we report the first observation in eukaryotic cells of toroid-shaped DNA objects, whose numbers vary dramatically across cells and species. Overall, our results show that dinoflagellate chromosomes exhibit both shared organizational features and pronounced species-specific deviations.

## INTRODUCTION

Dinoflagellates are a phylogenetically and ecologically diverse group of unicellular algae whose nuclear and chromosome morphologies are unique among eukaryotes. In fact, the dinoflagellate nucleus is so distinct that researchers coined a specific term for it: the dinokaryon [1]. Interphase chromosomes in dinoflagellates have several distinguishing features as observed by traditional Transmission Electron Microscopy (TEM) [2–12]. They are condensed and approximately cylindrical in shape. Chromosomes frequently show alternating high/low electron-density bands, with each band oriented perpendicular to the chromosome’s long axis and multiple bands stacked parallel to the chromosome’s long axis. Arch-shaped patterns are also common in oblique sections [2, 4]. Finally, the surface of dinoflagellate chromosomes is often ridged rather than smooth.

Traditional TEM is limited by a two-dimensional field of view that does not span the entire 3D chromosome, much less an entire nucleus or cell. Therefore, the precise number of chromosomes present, their 3D morphology, and the degree of structural variability among chromosomes within a single nucleus, across cells, and across species, has not yet been systematically and quantitatively compared. Three-dimensional volume electron microscopy would allow for the collection of larger sample sizes, necessary to infer general and robust conclusions about dinoflagellate chromosomes.

Around 2500 dinoflagellate species have been identified [13, 14]. This remarkable species diversity has motivated recent efforts to characterize dinoflagellate genomic diversity through the “100 Dinoflagellate Genome Project” [15]. However, an analogous inquiry into the degree of morphological diversity among dinoflagellate chromosomes is currently lacking.

Several lines of evidence suggest that chromosome morphology may be conserved across dinoflagellate species. There is strong indication that all dinoflagellates do not use histones to compact and organize their genomes, unlike all other known eukaryotes [16–21]. For example, nucleosomes are not visible in transmission electron micrographs of dinoflagellate chromatin spreads [16, 17], and micrococcal nuclease digestion of dinoflagellate chromatin does not produce the ladder of DNA bands that is expected from nucleosomal protection [18]. Histone transcripts are present in very low quantities across a wide range of species [20], but histone proteins are not detected by Western blot [18] or mass spectrometry [19]. Instead of histones, dinoflagellates have Dinoflagellate Viral Nucleoproteins (DVNPs), a unique set of highly expressed [20], basic, DNA-binding proteins that localize throughout chromosomes [18]. Given the central role of histones in chromosome organization in other eukaryotes [22], their extensive replacement by DVNPs in dinoflagellates suggests a common alternative chromosome morphology.

There are also notable distinct features between dinoflagellate species that may contribute to species-specific chromosome morphology. First, in contrast to DVNPs, Histone-like Proteins (HLPs) are present only in a subset of dinoflagellate species [23]. These proteins bind DNA [24], localize to the chromosome periphery [25, 26], and may actively organize chromatin [25, 26]. Two evolutionarily distinct HLP groups (HLP-I & HLP-II), found in separate dinoflagellate clades, were acquired during independent horizontal gene transfer events from different bacterial donors [23]. Second, genome size varies drastically across species, ranging from 0.13 Gb to 271.94 Gb [27, 28], which may affect the overall DNA density in the nucleus. Lastly, genes are arranged differently in different species. Many dinoflagellate genomes contain tandem repeat arrays [29–31], in which multiple copies of the same gene occur adjacently and are transcribed in the same direction. Tandem repeats are thought to influence chromosome morphology as their location correlates with the boundaries of Topologically Associating Domains (TADs) [32, 33], which are disrupted by transcriptional inhibitors [32, 33]. The abundance of tandem repeats varies across species which may result in species-specific chromosome morphology. For example, in *Symbiodinium microadriaticum*, 50% of genes exist in arrays of 9 or more co-oriented genes [32], while in *Fugacium kawagutii* tandem repeats are less common and shorter with only 2% of genes existing in arrays of 4 or more co-oriented genes [34]. It remains unclear whether these species-specific differences, or additional uncharacterized factors, result in distinct chromosome morphologies among dinoflagellate species.

In this study, we acquire three whole-nucleus Focused Ion Beam Scanning Electron Microscopy (FIB-SEM) datasets at 4 nm voxel size for each of four species— *Symbiodinium microadriaticum, Breviolum minutum, Fugacium kawagutii*, and *Crypthecodinium cohnii* —and compare these datasets with previously published image volumes from four additional species. Our results indicate that chromosomes in all species are neither physically confined nor spatially clustered in the nucleus. Using quantitative shape analysis methods, we find that dinoflagellate species share a principal component comprising 88% of the total chromosome shape variance. Despite these commonalities, species-specific exceptions to morphological trends are prominent, such as exceptions to left-handed helical organization, exceptions to isolated chromosomes, and variable abundances of rod-, crescent-, and toroid-shaped DNA objects. These findings indicate that dinoflagellate chromosomes share common morphological features while also exhibiting substantial species-specific variability.

## RESULTS

### 3D FIB-SEM and deep learning segmentation reveal diverse chromosome morphologies

We selected a diverse set of dinoflagellate species for imaging and analysis. First, we chose *Symbiodinium microadriaticum, Breviolum minutum*, and *Fugacium kawagutii* because they have relatively small genomes (1.10 Gb [36], 1.50 Gb [37], and 1.18 Gb [34], respectively), which have been previously sequenced and characterized. To complement these photosynthetic species, we also included *Crypthecodinium cohnii*, which is heterotrophic and has a larger estimated genome size (25 Gb [38]). Finally, we used existing FIB-SEM data from *Symbiodinium pilosum* [39], *Brandtodinium nutricula* [40], *Ensiculifera tyrrhenica* [41], and *Kareniaceae sp*. [42] due to their availability, high resolution, and inclusion of complete nuclei. Altogether, these eight species span four distinct taxonomic orders, allowing us to compare across distant lineages, as well as among more closely related taxa within the Suessiales order (Fig. S1). Table I provides a comprehensive overview of all dinoflagellate cells imaged and analyzed in this study, representing the largest collection of dinoflagellate FIB-SEM images assembled to date.

**TABLE I.** Summary of dinoflagellate cells imaged and/or analyzed in this study.

| Order | Species | Data Origin | Voxel Size [nm] | Sample Preparation | FIB-SEM | Cell ID |
| --- | --- | --- | --- | --- | --- | --- |
| Suessiales | *Symbiodinium microadriaticum* | This study | 4 | HPF/FS<sup>a</sup> | Zeiss Crossbeam 350 | 1, 2, 3 |
| Suessiales | *Symbiodinium pilosum* | [39] | 8 | HPF/FS | Zeiss Crossbeam 550 | 1, 2, 3 |
| Suessiales | *Breviolum minutum* | This study | 4 | HPF/FS | Zeiss Crossbeam 350 | 1, 2, 3 |
| Suessiales | *Fugacium kawagutii* | This study | 4 | HPF/FS | Zeiss Crossbeam 550 | 1, 2, 3 |
| Gonyaulacales | *Cryptocodinium cohnii* | This study | 4 | HPF/FS | Thermo Fisher Helios 5 Hydra CX | 1 |
| Gonyaulacales | *Cryptocodinium cohnii* | This study | 4 | HPF/FS | Zeiss Crossbeam 350 | 2, 3 |
| Peridiniales | *Brandtodinium nutricula* | [40] | 8 | HPF/FS | Zeiss Crossbeam 540 | 1, 2, 3 |
| Peridiniales | *Ensiculifera tyrrhenica* | [41] | 8 | HPF/FS | Zeiss Crossbeam 550 | 1 |
| Gymnodiniales | *Karenia* sp. | [42] | 8 | HPF/FS | Zeiss Crossbeam 550 | 1, 2 |

<sup>a</sup> High-Pressure Freezing/Freeze SubstitutionTABLE II. *F. kawagutii* skeleton regions

We imaged three cells for each of our four species: *S. microadriaticum, B. minutum, F. kawagutii*, and *C. cohnii*. We prepared samples by cryo-fixation with freeze substitution (Methods) to preserve native cellular ultrastructure [43–45], and to ensure that cross-species comparisons were not confounded by differences in sample preparation (Table I) [39–42]. Briefly (Fig. 1 a), (1) dinoflagellates were cultured in nutrient-enriched seawater; (2) cells were rapidly vitrified under high pressure and then freeze-substituted with an organic solvent containing heavy metal stains and fixatives; (3) samples were iteratively milled with a focused ion beam and imaged with a scanning electron beam; (4) successive cross-sections were digitally aligned to generate a 3D reconstruction of each cell nucleus; (5) a deep learning model was used to accelerate the segmentation of chromosomes, nucleoli, and the nuclear membrane. Example FIB-SEM cross-sections of cells, nuclei, and segmented chromosomes are presented in Fig. S2 and all FIB-SEM image stacks and segmentations are explorable in the ATLAS interactive online viewer (Data Availability). The ATLAS viewer runs in a web browser, requires no software installation, and loads within seconds by storing image data as spatially partitioned, multi-resolution tiles that are dynamically retrieved as needed for the current view [46]. Finally, segmented objects were visualized and subjected to morphological analysis in 3D.

**FIG. 1.**
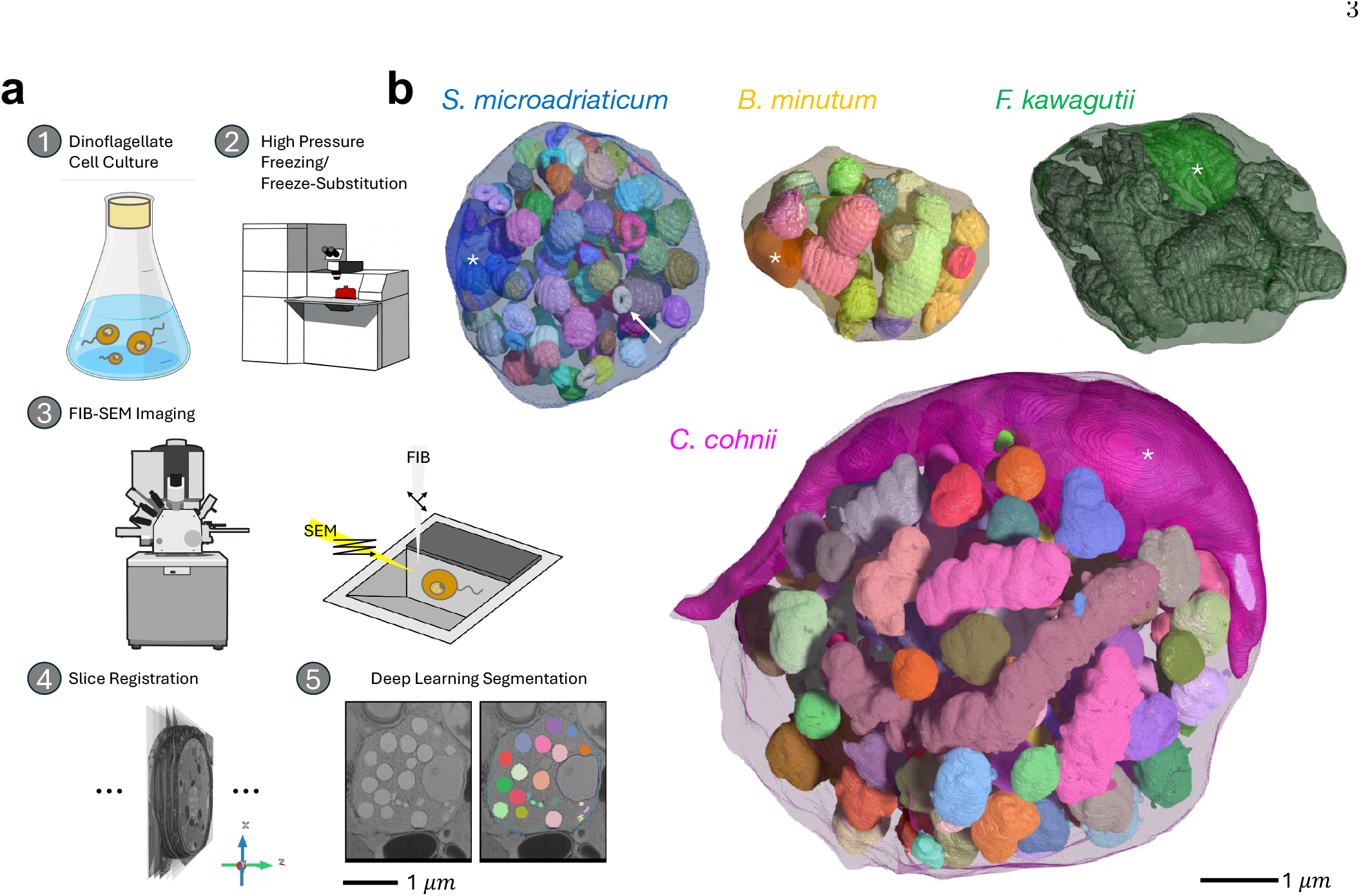
3D FIB-SEM and deep learning segmentation reveal diverse chromosome morphologies. a) Experimental workflow: 1. Individual dinoflagellate species were grown in artificial seawater with nutrients. 2. Dinoflagellate cells were rapidly vitrified under high pressure to prevent ice crystal formation. Frozen amorphous water was dissolved in an organic solvent with heavy metal stains and fixatives, then hardened and returned to room temperature, faithfully preserving cellular ultrastructure. 3. A cross-section of the cell was first exposed by an ion beam, then imaged with an electron beam. FIB-SEM image credit: [35]. 4. Successive cross-sections were digitally aligned to produce a 3D image. 5. A deep learning model was trained on select slices and used to dramatically accelerate segmentation of chromosomes, nucleoli, and the nuclear membrane. b) Example 3D reconstructions of dinoflagellate nuclei from each species imaged in this study, displayed to scale (Table I). Each chromosome is assigned a random color. Nucleoli are indicated with white asterisks. Example toroid is indicated with a white arrow.

Qualitatively, we observed substantial diversity in nuclear and chromosome morphology across dinoflagellate species. Figure 1 b shows 3D reconstructions of one representative nucleus from each species that we imaged: *S. microadriaticum, B. minutum, F. kawagutii*, and *C. cohnii*. Reconstructions of all nuclei, from all species, are presented in Fig. S3. *S. microadriaticum, B. minutum*, and *F. kawagutii* nuclei are comparable in size, while *C. cohnii* nuclei are roughly tenfold larger by volume. *C. cohnii* nucleoli are also larger and crescent-shaped, in contrast to the small, spherical nucleoli seen in the other species (Fig. 1 b, white asterisks). Chromosomes across all species are condensed and range from compact ellipsoids to elongated cylinders, with surface ridges visible on many, although *C. cohnii* chromosomes appear comparatively smoother. Chromosomes are discrete and individually isolated in most species, with the notable exception of *F. kawagutii*, whose chromosomes are inter-connected and form a single contiguous network. Finally, we observed toroid-shaped objects in several, but not all, nuclei. One striking example is *S. microadriaticum* Cell 3 which contains 14 distinct toroids (Fig. 1 b, white arrow). We next quantified each of these morphological features across all nuclei.

### Dinoflagellate chromosomes share unconfined, non-clustered organization in the nucleus

Dinoflagellates have been hypothesized to have highly condensed chromosomes in order to physically accommodate their large genomes (2.2-200 pg/nucleus compared to 3.2 pg/nucleus in humans [47]) within the limited nuclear volume [9, 48]. To evaluate whether dinoflagellate chromosomes are confined, we quantified the volume fraction of the nucleus occupied by chromosomes and nucleoli using our 3D segmentations. Surprisingly, we found that in all dinoflagellate species, regardless of genome size, chromosomes do not expand to fill the available nuclear space, occupying only 11-37% of the nuclear volume (Fig. 2 a). Nucleoli comprise an even smaller nuclear volume fraction, ranging from 1.5 to 9.3%. Nucleolar size, *V*_*No*_, increases with nuclear size, *V*_*Nu*_, according to a power-law, *V*_*No*_ ~ (*V*_*Nu*_)^0.88^ (Fig. S4 a), indicating that nucleoli occupy a smaller fraction of the nucleus as nuclear size increases.

**FIG. 2.**
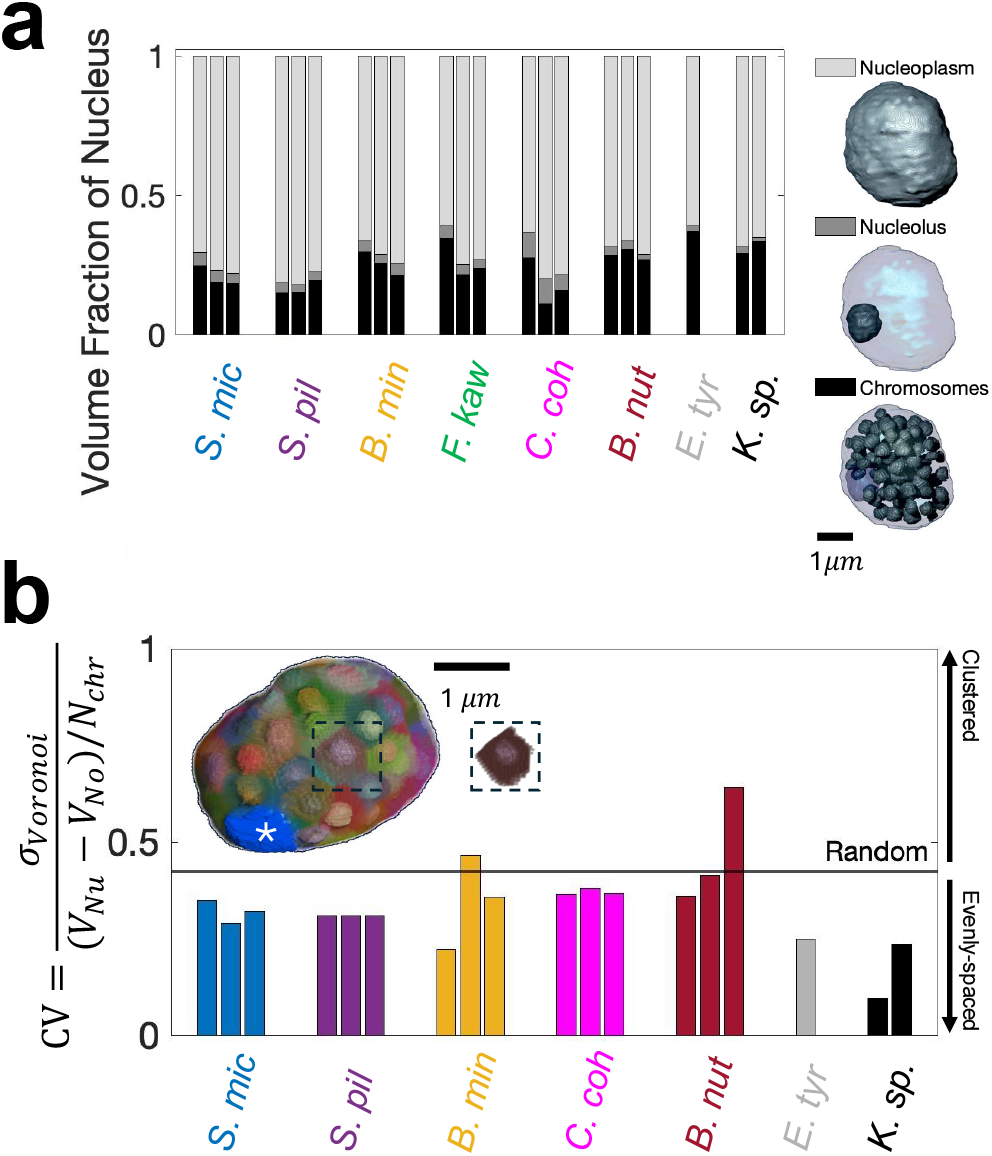
Dinoflagellate chromosomes share unconfined, non-clustered organization in the nucleus. Each bar is a cell. a) Volume fraction of the nucleus occupied by chromosomes (black), the nucleolus (dark grey), and nucleoplasm (light grey). b) Coefficient of variation (CV) of Voronoi volumes using chromosome centroids as seed points. An example nucleus with segmented chromosomes and Voronoi volumes is shown in the inset. An individual chromosome and its Voronoi volume is highlighted in the nucleus and displayed separately (dashed box). A white asterisk indicates the nucleolus. The CV numerator, *σ* _*V oronoi*_, quantifies the spread of the volume “claimed” by each chromosome, while the CV denominator, (*V*_*Nu*_ − *V*_*No*_)*/N*_*chr*_, is the average available space per chromosome. The horizontal solid line indicates the expected CV for randomly distributed points in 3D [49]. Values above/below the dashed line indicate more clustered/more evenly-spaced chromosomes, respectively.

We next investigated how dinoflagellate chromosomes are spatially distributed within the nucleus. As a first approximation, we used the convex hull of all chromosomes to estimate nuclear volume. This estimate closely agrees with the more precise volume calculated by segmenting the nuclear membrane (Fig. S4 b, c), corroborating the qualitative observation that chromosomes are located throughout the nucleus. To more precisely characterize how chromosomes are distributed, we quantified the heterogeneity of chromosomal positions using a Voronoi simulation [50]. Chromosome centroids were taken as seed points which concurrently dilated in 3 dimensions until they contacted each other, a nucleolus, or the nuclear membrane (Methods). The resulting Voronoi volumes (Fig. 2 b, inset) represent the volume “claimed” by each chromosome. If chromosomes are clustered, then the standard deviation of the Voronoi volumes, *σ* _*V oronoi*_, would be high relative to the average available space per chromosome, (*V*_*Nu*_ − *V*_*No*_)*/N*_*chr*_ [50]. Alternatively, if chromosomes are evenly spaced, *σ* _*V oronoi*_ would be relatively small. Regardless of species or cell, the Coefficient of Variation (CV) of Voronoi volumes is below or near what is expected for a random distribution of points in 3D [49] (Fig. 2 b, horizontal dashed line) indicating that dinoflagellate chromosomes are not clustered spatially. Together, these findings establish two common organizational features, namely that dinoflagellate chromosomes are consistently unconfined and spatially distributed throughout the nucleus.

### Dinoflagellate chromosomes exhibit diverse shapes, with similar shape variational components across species and cells

Our 3D segmentations revealed extensive morphological diversity among objects within the nucleus. We observed many shapes including rods, crescents, toroids, small and large “blobs” with smooth surfaces, ellipsoids with surface ridges, irregular shapes, half-helical and half-disordered chromosomes, T-junctions, and large contiguous networks (Fig. 3 a). To quantitatively assess these morphological differences, including their prevalence and distribution across species and cells, we adapted an approach previously developed to analyze human cell and nuclear shape [51]. We used Spherical Harmonic Expansion (SHE) to parametrize each chromosome’s shape and Principal Component Analysis (PCA) to visualize the resulting shape landscape and identify any dominant modes of shape variation [51] (Methods). First, chromosomes were re-oriented so their long axes point vertically, removing any orientation dependence from the shape analysis (Fig. S5 a). Then, each chromosome was parametrized by a set of spherical harmonic functions (Fig. S5 a). The error between the segmented shape and the SHE reconstruction was calculated per chromosome and chromosomes with a reconstruction error below a threshold were included in our analysis (Methods).

**FIG. 3.**
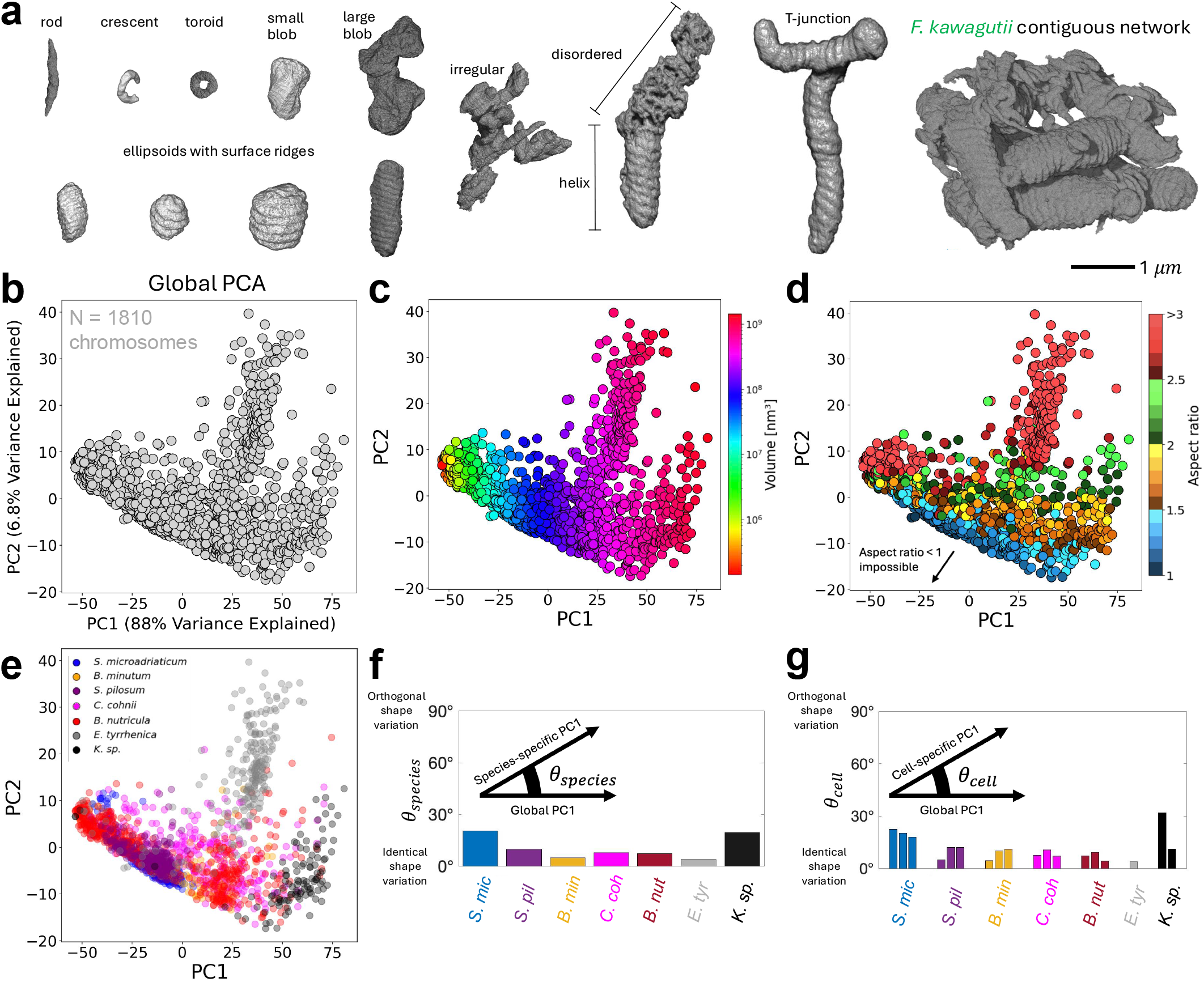
Dinoflagellate chromosomes exhibit diverse shapes, with similar shape variational components across species and cells. a) To-scale examples of diverse DNA object morphologies found in dinoflagellate nuclei. b) Principal Component Analysis (PCA) after Spherical Harmonics Expansion (SHE) [51] identifies shape modes that capture significant (*PC*1 = 88%, *PC*2 = 6.8%) chromosome shape variation across all dinoflagellates (*N* = 1810 chromosomes). c) PC1 is highly correlated (*R*^2^ = 0.82) with chromosome volume. d) PC2 is moderately correlated (*R*^2^ = 0.44) with chromosome aspect ratio. e) Some species occupy distinct regions in chromosome shape-space. f) Angle between global *PC*1 (same as in b) and species-specific *PC*1 where PCA and SHE only use chromosomes from a single species. g) Angle between global *PC*1 (same as in b) and cell-specific *PC*1 where PCA and SHE only use chromosomes from a single cell. Each bar is a cell.

Altogether, we parametrized *N* = 1810 (85% of all) chromosomes across 7 species (Methods) and used PCA to identify the two orthogonal axes that explain the greatest amount of shape variation (Fig. 3 b). We found that the first principal component, *PC*1, accounts for a surprisingly large fraction (88%) of the total shape variance, while the second principal component accounts for nearly all the remaining variance (6.8%). The compression of most shape variation into a low-dimensional representation reveals that dinoflagellate chromosome diversity is surprisingly parsable. By contrast, the top eight principal components describing human cell and nuclear shape variation account for less variance (69.4%) than our *PC*1 alone [51].

To interpret what shape variation each principal component captures, we sampled along the *PC*2 = 0 and *PC*1 = 0 lines, respectively, and visualized those resulting shapes. We observed approximately isotropic growth while increasing the *PC*1 coordinate, and growth along one axis while increasing the *PC*2 coordinate (Fig. S5 b). Indeed, *PC*1 highly correlates with chromosome volume (*R*^2^ = 0.82, Fig. 3 c) and *PC*2 moderately correlates with aspect ratio (*R*^2^ = 0.44, Fig. 3 d). The bottom-left corner of the PCA plot is empty because a chromosome’s aspect ratio (i.e. height/width) cannot be less than 1, given their imposed vertical orientation.

Next, we compared the distribution of chromosomes in shape space across species. We observed that some species (*S. microadriaticum & S. pilosum*) occupy overlapping regions in shape space, while other species (*K. sp & E. tyrrhenica*) occupy more unique regions of shape space (Fig. 3 e). Repeating dimensionality reduction using an equal number of randomly selected chromosomes from each species resulted in a near identical projection with the angle between this *PC*1 and the global *PC*1 equal to 1.4° (Fig. S5 c), confirming that the PCA is robust to differences in chromosome number across species, and to differences in the number of cells imaged per species (Table I).

Given that some species occupy distinct regions of shape space, we sought to quantify chromosome shape variation for each species separately and compare them to the global *PC*1 axis (horizontal axis, Fig. 3 b-e). To do so, we repeated the PCA using only data from one species (e.g. Fig. S5 d) and computed the angle *θ*_*species*_ between the species-specific *PC*1 and the global *PC*1 (Fig. 3 f). A small angle *θ*_*species*_ ≈0° indicates that the variation of chromosome shape within a particular species (species-specific *PC*1) is similar to that across all species (global *PC*1), while a large angle *θ*_*species*_ ≈90° indicates that a species’ chromosome shape variation is distinct from the global trend. We found that *θ*_*species*_ < 21° for all 7 species with some species having even smaller angles (Fig. 3 f). We also found the standard deviation of the angles to be 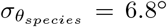. Therefore, dinoflagellate chromosomes share organizational features across the diverse species analyzed. We also considered cell-to-cell variation within each species, repeating the PCA using only data from a single cell and analogously computing the angle *θ*_*cell*_ between the cell-specific *PC*1 and the global *PC*1. We find that *θ*_*cell*_ remains small (< 32°) and is highly similar within each species, with an average intra-species standard deviation of 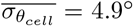 (Fig. 3 g). One notable outlier is *K. sp*. where chromosomes in one nucleus appear longer and more bent than in the other nucleus (Fig. S3). However, overall we observe slightly greater inter-species variation than intra-species variation as 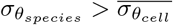.

We further characterized dinoflagellate chromosome shape by inspecting their cross-sectional profiles, allowing for quantification of differences in curvature at the chromosome ends. We approximated the chromosome’s cross-section using a rectellipse curve, 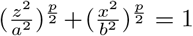, which has the chromosome length, 2*a*, the chromosome width, 2*b*, and the curvature, *p*, as flexible parameters (Fig. S6 a). Large curvature values, *p* ≥10, correspond to minimal taper at the chromosome ends, while *p* = 1 corresponds to maximum taper. Cross-sectional profiles of segmented chromosomes from *S. microadriaticum, S. pilosum, B. minutum*, and *E. tyrrhenica* are more consistent across chromosomes, while those from *C. cohnii* and *K. sp*. are more varied, with less overlap among profiles within each species (Fig. S6 b). Some *C. cohnii* and *K. sp*. chromosomes have profiles that deviate from the rectellipse curve, exhibiting irregular rather than smoothly arced boundaries. Interestingly, cross-sectional chromosome profiles from *B. nutricula* segregate into two populations, one for smaller chromosomes and one for larger chromosomes (Fig. S6 b). Consistent with this, we found two distinct clusters of *B. nutricula* chromosomes along *PC*1 (~ volume axis) in the species-specific PCA shape space (Fig. S5 d). In general, we found that dinoflagellate chromosomes are significantly tapered 1.7 ≤ *p* ≤ 2.7, independently of their aspect-ratio (Fig. S6 c). Therefore, we conclude that tapered ends are a prevalent shape characteristic among many, but not all, dinoflagellate chromosomes.

Both our SHE/PCA and our cross-sectional profile analysis indicate that the shapes of dinoflagellate chromosomes have several common features, yet occasional species-specific, cell-specific, or chromosome-specific deviations preclude a fully universal description.

### Many, but not all, dinoflagellate chromosomes are left-handed helices

Some dinoflagellate chromosomes have ridges on their surface. We provide example images of chromosomes with and without surface ridges in the inset of Figure 4 a. Detection of ridges requires sufficiently high resolution (≪ 40 nm voxel size, depending on ridge spacing) (Fig. S7 a). Using our high-resolution FIB-SEM images and others from the literature (Table I), we found that 44% of chromosomes across all species have surface ridges. However, the prevalence of surface ridges varies across species; for example, only 11% of *C. cohnii* chromosomes have ridges while 91% of *E. tyrrhenica* chromosomes have ridges (Fig. 4 a). For chromosomes that have ridges, we manually annotated the location of each ridge using Fiji’s drawing tools [52] (Fig. S7 b) and found that the number of ridges increases linearly with chromosome length (*R*^2^ = 0.836, Fig. 4 b). This suggests that a ridge constitutes a repeatable structural unit. The slope of the best-fit line gives the average ridge spacing and is 6.1 ridges*/µ*m or 160 nm per ridge. *S. pilosum* chromosomes have the smallest average ridge spacing of 73 nm, while *C. cohnii* chromosomes have the largest ridge spacing at 206 nm. Our measurements are consistent with recent chromosome band spacing measurements performed on conventionally fixed, expanded, dinoflagellate cells — 160 nm *±* 21 nm in *K. papilionaceae* and 190 nm *±* 24 nm in *Prorocentrum sp*. [53].

**FIG. 4.**
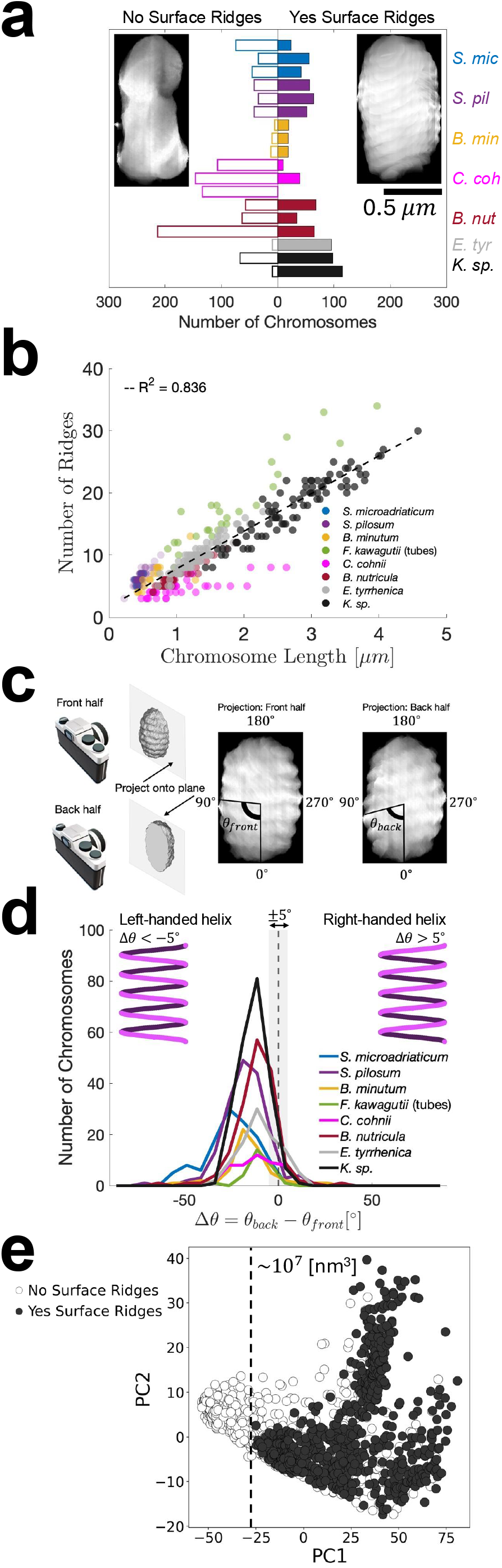
Many, but not all, dinoflagellate chromosomes are left-handed helices. a) Number of chromosomes without surface ridges (left, open bars) or with surface ridges (right, closed bars). Each bar is a cell. Example chromosomes are shown as insets. b) Chromosome length versus number of surface ridges. Each data point is a chromosome, except for *F. kawagutii*, where each data point is a tube (see Fig. S7 b). A linear relationship (*R*^2^ = 0.836) indicates there is a repeatable organizational unit in many dinoflagellate chromosomes. c) Schematic showing ridge angle extraction. Each chromosome is projected inwards from the front, or the back, onto its bisecting plane, and viewed from the same perspective (camera). The angle of ridges relative to the vertical axis, *θ*, is estimated for each half. d) Δ*θ* distinguishes left-handed helices from right-handed helices (see also Fig. S8). Most ridged dinoflagellate chromosomes are left-handed helices. Grey region indicates |Δ*θ*| < 5°. e) Surface ridges are present only for chromosomes larger than ~ 10^7^ nm^3^. Here, the *PC*1, *PC*2 shape axes are the same as in Fig. 3.

In addition to ridge spacing, analyzing ridge orientation can reveal whether chromosomes are helically organized and, if so, the handedness of the helix. Our approach builds on work from Oakley & Dodge [54], who tracked ridge orientation across serial TEM sections of a single *Glenodinium hallii* chromosome, and De La Tour & Laemmli [55], who used optical sectioning to reveal a helical topoisomerase II scaffold in human metaphase chromosomes. To quantify ridge orientation, we first enclosed isolated chromosomes in a bounding box and re-oriented them so that their long axes were aligned vertically (Methods, Fig. S5 a). Using simulated 3D binary images of helices with known handedness, we verified this procedure did not alter the chirality of the helix. For *F. kawagutii*, which does not have isolated chromosomes, we placed a cylindrical cropping region around each thick “tube” and re-oriented that segment vertically (Fig. S7 c). We also used these “tubes” to determine linear ridge density in Fig. 4 b. Next, we divided each chromosome into a front and back half, and projected each hemi-volume inwards onto the mid-plane (Fig. 4 c). From a single perspective (e.g. the “front”), we manually quantified the angle of these ridges relative to the chromosome long axis using Fiji’s protractor tool [52]. Verification of ridge angle measurements was done via visual inspection after superimposing an angle indicator onto the projected chromosome halves (Fig. S7 d). All angle measurements are available as a series of galleries, with each gallery corresponding to chromosomes in a single cell (Supplementary Data S1-S18). The difference of the back-half surface ridge angle, *θ*_*back*_, and the front-half surface ridge angle, *θ*_*front*_, Δ*θ* = *θ*_*back*_ − *θ*_*front*_ can be used to distinguish left-handed (Δ*θ* < 0°) and right-handed (Δ*θ >* 0°) helices, as well as flat discs (Δ*θ* = 0°) with arbitrary tilt (Fig. S8). We calculated Δ*θ* for every chromosome with surface ridges and plot the distributions for each species in Fig. 4 d. As Δ*θ* = 0 is an infinitely thin domain of the distribution, we considered a *±* 5° tolerance to account for uncertainties in angle extraction (Fig. 4 d, S7 d, grey regions). We found that for all species, the Δ*θ* distribution contains a single peak at a negative value, indicating that many dinoflagellate chromosomes are left-handed helices (Fig. 4 d). Specifically, 82% of all chromosomes with ridges are left-handed (Δ*θ* < 5°), while only 15% of chromosomes with ridges have flat discs (|Δ*θ*| < − 5°). Exceptionally, 3.1% of chromosomes with ridges are right-handed (Δ*θ >* 5°), establishing that both left- and right-handed helical chromosomes exist in dinoflagellate nuclei.

Finally, we examined whether the presence or handedness of helicity correlates with chromosomal shape features. We found overlap in the regions of shape space occupied by chromosomes with and without surface ridges (Fig. 4 e). Nevertheless, there is a minimum size threshold of ~ 10^7^ nm^3^ below which no ridges were observed (Fig. 4 e). We also found significant overlap in the PCA shape space regions occupied by left-handed helices (Δ*θ* < − 5°) and flat discs (| Δ*θ* | < 5°), while right-handed helices (Δ*θ >* 5°) appear to have small aspect ratios (Fig. S9).

Together, these findings demonstrate that while certain organizational features — such as ridge number scaling with chromosome length and left-handed helical ridge arrangement — are broadly conserved, the extent of surface ridge organization varies considerably across individual chromosomes and across species.

### One contiguous chromosome network distinguishes

#### *F. kawagutii* from other species

Compared to other species, *F. kawagutii* chromosomes exhibit a qualitatively different organization. Indeed, *F. kawagutii* nuclei contain large cylindrically shaped “tubes” with left-handed helical surface ridges (Fig. 4 d) that are interconnected by many small “bridges”, forming a contiguous network (Fig. 5 a, Supplementary Videos S1-S3).

**FIG. 5.**
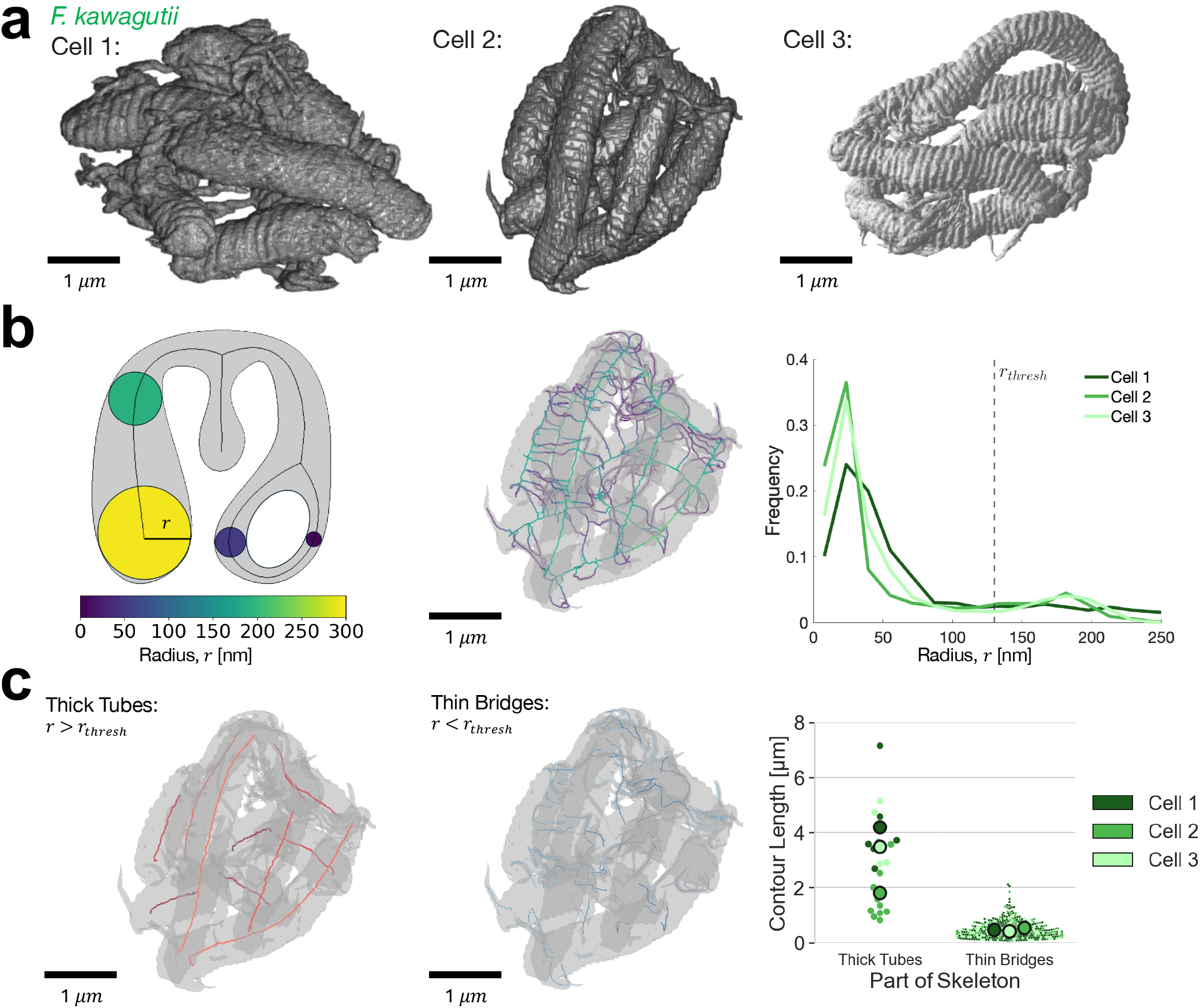
One contiguous chromosome network distinguishes *F. kawagutii* from other species. a) Segmented *F. kawagutii* chromosomes show one contiguous network in each of three cells. b) Left panel: Skeletonization schematic (modified with permission from [56]) showing radii, *r*, of the largest inscribed spheres centred at points along the skeleton. Middle panel: Visualization of chromosome skeleton colored by radius. Right panel: A histogram of radii sampled uniformly along the skeleton informed a threshold choice of *r*_*thresh*_ = 130 nm separating thick from thin regions. c) Visualization of skeleton after application of radius threshold. Thick tubes (*r > r*_*thresh*_, left) are separated from thin bridges (*r* < *r*_*thresh*_, right). Each thick tube and thin bridge is assigned a random color shade for better distinguishability. See Fig. S10 for other cells. Contour lengths of each thick tube and thin bridge, colored by cell.

Skeletonization is a process that reduces a complex 3D object to its central core, or medial axis, while preserving the object’s topology and branching structure. Medial axis skeletonization starts by measuring how far each voxel is from the object’s surface, and then extracts the set of points that is locally farthest from the boundary, reducing the object to a simplified, thin representation. To characterize the *F. kawagutii* chromosome network, we applied the Erosion Thickness skeletonization algorithm (Methods) [57, 58], which offers two key advantages over other methods. First, it skeletonizes thin “bridges” while remaining insensitive to similarly sized surface ridges, and second, it colors the skeleton by the local maximally inscribed sphere radius, *r*, providing a measure of spatial thickness at each point (Fig. 5 b). Roughly 35000 overlapping spheres are located 5 nm apart along the skeleton contour. We used the histogram of these spheres’ radii to define a threshold, *r*_*thresh*_ = 130 nm, that distinguishes thick “tubes” from thin “bridges” (Fig. 5 b, Methods). In Fig. 5 c, we show an example of the thresholded skeleton for Cell 2; skeletons for Cells 1 and 3 are provided in Fig. S10. We counted 6-11 “tubes” and 121-216 “bridges”, depending on the cell (Table II). We next measured the contour lengths of each “tube” and “bridge” and found that “tubes” range from 0.5-7.5 *µ*m in length, while “bridges” are typically < 1 *µ*m (Fig. 5 c). Cell 2 has more “tubes” compared to Cells 1 & 3, but they are also shorter. The total volume of the entire chromosome network is similar across cells (Table II), which may suggest that DNA content can be redistributed between “tubes” and “bridges”. Overall, *F. kawagutii* stands as a striking exception among dinoflagellates: unlike other species, whose chromosomes are isolated and discrete, its chromosomes adopt a uniquely networked organization.

**TABLE II.**
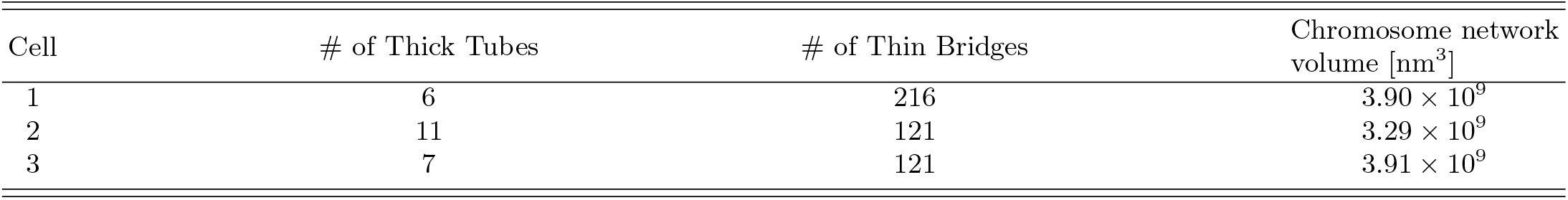
*F. kawagutii* skeleton regions.

### Dinoflagellate nuclei contain toroidal DNA objects with differing abundances across cells and species

In addition to the *F. kawagutii* chromosome network, we observed other unexpected structures in dinoflagellate nuclei such as rods, crescents, and toroids. The abundances of these objects vary substantially across species and cells (Fig. 6 a). For example, rods are highly abundant in *S. microadriaticum, C. cohnii*, and *B. nutricula* but totally absent in *S. pilosum*. Many toroids were observed in *S. microadriaticum*, while most other species had at least one, except for *F. kawagutii* and *C. cohnii*, where no toroids were observed. Remarkably, in *S. microadriaticum*, Cell 3 contains 14 toroids, whereas Cell 1 contains only 2. We measured the dimensions of each object and found that rods, crescents, and toroids have comparable widths, ranging from 20 to 200 nm. However, rods are typically shorter (~ 500 nm) than crescents (300 − 1600 nm) and toroids (800 − 2400 nm) (Fig. 6 b). For context, rods/crescents/toroids have roughly 40-6 fold smaller volume compared to chromosomes.

**FIG. 6.**
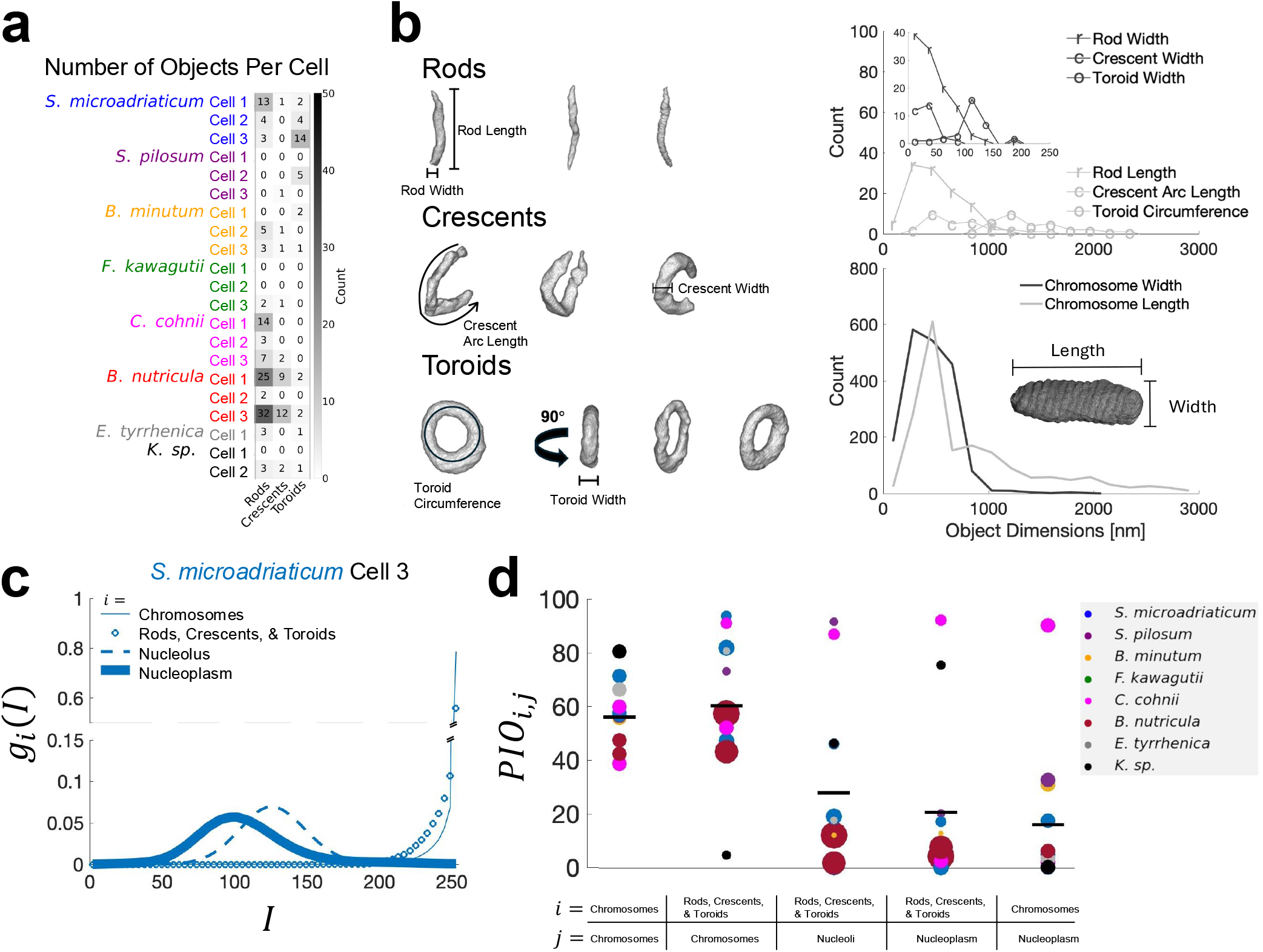
Dinoflagellate nuclei contain toroidal DNA objects with differing abundances across cells and species. a) Count of rods, crescents, and toroids in each cell. b) Rod, crescent, and toroid dimensions compared to chromosome dimensions. c) FIB-SEM voxel intensity distribution *g*_*i*_(*I*) for each segmented object type *i*. The distribution for rods/crescents/toroids overlaps significantly with that for chromosomes. d) Percent intensity overlap *PIO*_*i,j*_ of intensity distributions *g*_*i*_(*I*) and *g*_*j*_ (*I*) between object types *i* and *j* (Methods). *PIO* quantifies the similarity between two object type’s electronic properties and affinity for heavy metal stains. Positive control comparing random pairs of chromosomes; negative control comparing chromosomes to nucleoplasm. Rods/crescents/toroids compared to chromosomes, nucleoli, and nucleoplasm. Each dot is a cell. Dot size indicates object sample size. Color indicates species. Black lines indicate mean, 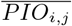, across all cells holding object types *i* and *j* fixed.

We hypothesized that these objects may represent chromosome fragments or extrachromosomal DNA. To determine whether these structures exhibit compositional similarities to chromosomes, we analyzed their FIB-SEM voxel intensity distributions, *g*(*I*). Voxel intensity is determined by the local composition of the sample, including its electronic properties and affinity for heavy metal stains. We compared the FIB-SEM voxel intensity distributions of rods/crescents/toroids with those of chromosomes, the nucleolus, and the nucleoplasm (Methods). For example, in *S. microadriaticum* Cell 3 we found the intensity distribution of rods/crescents/toroids to be qualitatively similar to that of chromosomes and distinct from the nucleolar and nucleoplasm distributions (Fig. 6 c). To quantify this similarity, we computed the Percent Intensity Overlap, *PIO*, (Methods) between the voxel intensity distributions for every dinoflagellate nucleus containing rods, crescents, or toroids. As a positive control, we first computed the *PIO* between random chromosome pairs and found 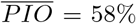 (Fig. 6 d). As a negative control, we computed *PIO* between chromosomes and the nucleoplasm and found 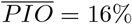. We then calculated *PIO* for rods, crescents, and toroids against chromosomes 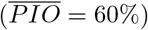, the nucleolus 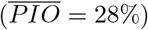 and the nucleoplasm 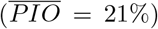. These results suggest that rods/crescents/toroids are compositionally similar to chromosomes and may therefore contain DNA. Ultimately, the pronounced variability in rod, crescent, and toroid abundances across cells and species underscores the substantial cell-to-cell heterogeneity of dinoflagellate nuclear organization.

## DISCUSSION

Using 3D FIB-SEM imaging, we examined the chromosome morphology of several dinoflagellate species, revealing both common features and species-specific differences. Commonalities include a consistent lack of confinement or spatial clustering of chromosomes within the nucleus (Fig. 2). Furthermore, nearly all dinoflagellate chromosome shape variation (95%) is explained by two principal components that correlate with chromosome volume and aspect ratio (Fig. 3 b-d). Despite these commonalities, we also observed several notable exceptions. For instance, only 44% of dinoflagellate chromosomes have surface ridges (Fig. 4 a). When surface ridges are present, they generally correspond to left-handed helices; however, 15% of chromosomes with ridges have flat discs and 3.1% of chromosomes with ridges are right-handed helices (Fig. 4 d). *F. kawagutii* differs from other species in that its chromosomes are not discrete well-separated entities, but instead connected into a single contiguous network (Fig. 5). Finally, rod-, crescent-, and toroid-shaped DNA objects are present in most dinoflagellate nuclei, but their abundances vary dramatically across cells and species (Fig. 6 a).

Surprisingly, we found that chromosome number is variable — even within a single species (Table III). This variation may reflect ongoing DNA replication during S phase, spatial overlap of multiple chromosomes, or possibly an inherent unstable karyotype. Some species have low variation, such as *S. microadriaticum, S. pilosum*, and *B. minutum*, while the greatest variation was observed in *B. nutricula*, where Cells 2 & 3 differ in chromosome number by almost 3-fold. Complicating this picture, chromosome counts also vary depending on the method used, such as between FIB-SEM, light microscopy, and high-throughput chromosome conformation capture (Hi-C) [21, 32–34, 59, 60] (Table III). For *S. microadriaticum*, estimates of chromosome number from all three methods are in close agreement. By contrast, *B. minutum* and *F. kawagutii* chromosome counts show substantial disagreement across methods (Table III). The large number of *B. minutum* chromosomes predicted by Hi-C may be due to an incomplete genome assembly, which covers only 39% of the estimated genome size [33, 61]. The Hi-C estimate for *F. kawagutii* – 99 chromosomes [21, 61] – is also much greater than those from both electron and light microscopy. Indeed, we could not distinguish individual chromosomes in *F. kawagutii* using FIB-SEM and only 7–11 chromosomes were observed by confocal imaging [34, 59, 60]. Interestingly, these confocal estimates are consistent with the number of thick “tubes” that we observed by FIB-SEM (Table II). Nevertheless, our discovery of thin “bridges” that connect these tubes (Fig. 5 c) reveal that “tubes”, and likely the DAPI-stained objects, are not isolated chromosomes. Thus, the true number of *F. kawagutii* chromosomes remains undetermined.

**TABLE III.**
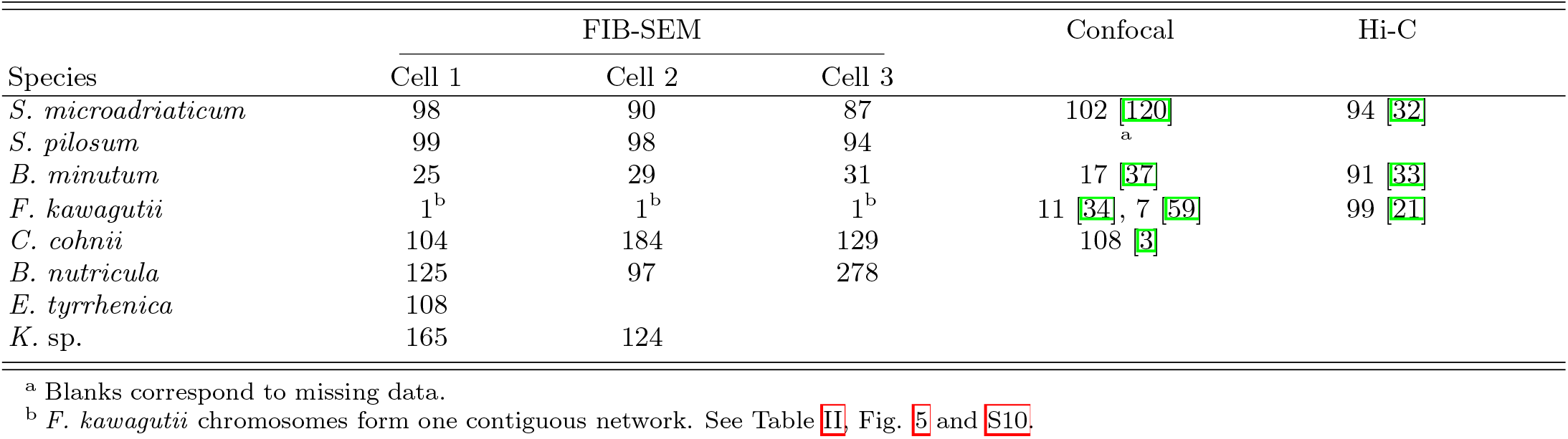
Chromosome counts by method.

We also note that *B. minutum* chromosomes are on average 3.3-fold less numerous but 3.4-fold larger than more distal dinoflagellate species, *S. microadriaticum* and *S. pilosum* (Fig. S1), which may indicate past chromosome fission events.

Similar to chromosome number, the structure of individual dinoflagellate chromosomes has also been difficult to determine. The most prominent model invoked in the literature is a cholesteric liquid crystal (CLC), which consists of rotating stacks of flat discs [2, 4–6, 15, 62, 63].

However, our recent Hi-C-constrained polymer simulations found no evidence for such organization, instead supporting more irregular chromosomal conformations [61]. Our FIB-SEM data corroborate this non-CLC view: 56% of all dinoflagellate chromosomes lack surface ridges entirely — inconsistent with a layered architecture — and only 6.4% of all dinoflagellate chromosomes have surface ridge angle measurements consistent with flat discs (|Δ*θ*| < 5°) (Fig. 4 d). Instead, our data reveal that a substantial fraction of dinoflagellate chromosomes are helical.

A key advantage of FIB-SEM over Hi-C is its ability to resolve helix handedness [64, 65]. Prior inferences of dinoflagellate chromosome handedness have relied on circular dichroism [66], *ex-vivo* approaches [67], or 2D image data [68] — each limited to a small number of chromosomes in a single species. These studies reported both left-[66, 67] and right-handed [68] chromosomes, consistent with our observations (Fig. 4 d). By directly imaging all chromosomes across many nuclei and multiple species, we can draw broader conclusions: left-handed helices are relatively common (36%), and right-handed helices are exceptionally rare (1.4%). What earlier work did not emphasize or quantify is that most dinoflagellate chromosomes (56%) lack ridges entirely.

The prevalence of chromosomes lacking surface ridges may partly reflect dynamic, transcription-driven remodeling. For example, the half-helical, half-disordered chromosome in Fig. 3 a) spans the interface of the nucleolus, with the disordered half located inside the nucleolus and the helical half in the nucleoplasm. This arrangement suggests that ribosomal DNA transcription locally disrupts chromosome structure. Similar nucleolus-associated loss of structure has been reported in other dinoflagellate species [40, 69], suggesting this may be a general phenomenon.

A surprising observation is the existence of rods, crescents, and toroids in dinoflagellate nuclei (Fig. 6). The high overlap of intensity profiles of these objects to that of chromosomes implies that they have similar affinity for heavy metal stains (Fig. 6 c, d). Rods, crescents, and toroids thus appear compositionally similar to chromosomes, suggesting they too contain DNA. DNA toroids, while unexpected, are not exclusive to dinoflagellates. They have also been observed in bacterial cells under starvation stress [70], in viral capsids [71], *ex vivo* in treated mouse sperm nuclear lysate [72], and *in vitro* [73–79]. To our knowledge, however, this is the first observation of DNA toroids inside intact eukaryotic cells.

DNA rods and toroids have been shown to dynamically interconvert *in vitro* [80, 81] as well as *in silico* [82, 83]. The comparable widths of rods, crescents, and toroids, and the comparable arc-length of crescents to the circumference of toroids (Fig. 6 b), suggest that these structures may similarly interconvert *in vivo*. Rod, crescent, and toroid widths are also comparable to the size of surface ridges (73-210 nm), raising the possibility that they share the same underlying organization.

Alternatively, instead of rods bending into crescents and bending further into toroids, another potential explanation for toroidal shape may be the inherent topological constraint imposed by circular DNA. DNA minicircles are small (0.8-6 kb), gene-encoding, circular DNA molecules present in dinoflagellates [84–87]. Using Fluorescent *In Situ* Hybridization (FISH), DNA minicircles have been localized to dinoflagellate plastids [86] and nuclei [85], though sequence-based tagging and high resolution microscopy have not yet been combined to conclusively determine if minicircles form 3D toroid shapes. However, minicircle lengths do predict a range of arc-lengths/circumferences (0.8-6 kb × 0.34 nm/bp [88] = 272-2040 nm) consistent with our measurements for crescents and toroids (Fig. 6 b). Furthermore, minicircle copy number can vary by more than an order of magnitude [87, 89] in the same species, consistent with our observations of highly variable toroid abundances across cells.

Regardless of whether toroids arise from linear or circular DNA, toroid formation requires a condensing agent to neutralize negatively charged DNA. Polycationic compounds are capable of inducing toroid formation *in vitro* at sufficiently high concentrations [73–79]. Dinoflagellate nucleoplasm contains several such candidates: high concentrations of divalent cations such as Mg^2+^ and Ca^2+^ [90], DVNPs [18], and HLPs [23]. Both DVNPs and HLPs generally have basic isoelectric points [20], and are known to bind DNA and to localize with chromosomes in cells [18, 25, 26].

HLPs are highly abundant in the nucleus of some dinoflagellate species [24–26, 91–94], and bacterial Heat Unstable (HU) proteins — the closest orthologs to HLPs in dinoflagellates [23] — share high sequence homology with them [95]. Given that increasing HU concentration *in vitro* resulted in a higher occurrence of rods compared to toroids [96], cell-to-cell variation in HLP concentration may explain why rod and toroid proportions differ across dinoflagellate cells.

In addition to toroids, poly-cationic compounds may also contribute to the highly condensed state of dinoflagellate chromosomes, which remain condensed throughout interphase [8, 97, 98]. Electrostatic interactions may provide an alternative force driving chromosome condensation as dinoflagellate chromosomes are not confined, occupying only 20–40% of the nuclear volume (Fig. 2 a). Our findings are consistent with prior TEM stereology measurements of chromosomal nuclear volume occupancy: 46% in *P. micans* and 36% in *P. triestinum* [67]. Dinoflagellate chromosomes are thus not space-filling — a marked contrast to human nuclei, where DNA is distributed nearly everywhere in the nucleoplasm [22, 99, 100]. A lack of confinement further contrasts dinoflagellate chromosomes with sperm chromosomes, which adopt similarly condensed and twisted structures, yet are tightly confined by the nuclear envelope [101].

Table S1 summarizes key genomic and morphological features of dinoflagellate chromosomes across species and orders, comparing observations from our 3D FIB-SEM analysis with those from the literature. Across orders, a condensed morphological (“dinokaryonic”) phenotype — characterized by cylindrical-shaped chromosomes with bands of alternating intensity, surface ridges, and arches visible in cross-section — is common across non-parasitic dinoflagellates. This phenotype is observed in species bearing either HLP-I or HLP-II variants [9, 23]. Our 3D analysis revealed that *S. microadriaticum* and *B. minutum* share similar chromosome morphologies despite their drastically different prevalence of tandem repeats [32, 34, 37], suggesting that large-scale 3D chromosome architecture may be conserved independently of sequence-level genomic organization. We also discovered that *F. kawagutii* is unique in its contiguous network morphology. TEM studies show additional variation within the Suessiales order: chromosomal arches in cross-section are present in some species but absent in others, and surface ridges were not observed in *B. andersenii* [102]. Several species within the order Gymnodiniales also lack surface ridges: many *Testudoinium* species [103, 104], as well as *P. jejuensis* [105].

Parasitic dinoflagellate orders (Table S1) — Syndiniales, Noctilucales, and Blastodiniales — are especially exceptional in their chromosome morphologies. For example, *A. ceratii* completely lack dinokaryonic chromosomes [106], *B. sp*. lack dinokaryonic chromosomes while feeding inside the host as a trophont but only sometimes possess chromosomes with surface ridges as a spore [107], and *N. scintillans* lack dinokaryonic chromosomes as a trophont but possess fully dinokaryonic chromosomes as a spore [108]. These observations suggest a continuum of chromosome morphologies within dinoflagellate parasites from more traditional eukaryotic-appearing to more dinokaryonic. *N. scintillans* is particularly interesting because it expresses both DVNP and HLP-II, which have been hypothesized to contribute to dinokaryonic morphology in non-parasites [18, 23]. However, it remains unknown whether expression of these proteins is life-stage-dependent or whether they result in expression-level-dependent chromosome morphology. *A. ceratii* has the smallest dinoflagellate genome (0.13 Gb [27]) and dinokaryonic features have not been reported, consistent with our observation that a minimum chromosome size is required for surface ridge formation (Fig. 4 e). *P. balticum* [109], *D. kwazulunatalensis*, and *D. capensis* [110] are “dinotoms”, which have two nuclei per cell: a dinokaryon nucleus with condensed chromosomes containing surface ridges, and a traditional eukaryotic nucleus lacking condensed chromosomes. Thus, chromosome morphology can be a function not only of life cycle but also of nuclear identity, with multiple distinct morphologies coexisting within the same cell. Overall, our FIB-SEM imaging, together with prior TEM observations, reveals a rich and heterogeneous chromosomal landscape that is not fully captured by the prevailing CLC model.

Where earlier studies were constrained to population-averaged sequence-based methods [21, 32, 33, 61] or limited field of view 2D imaging [2–12], we now have a three-dimensional single-cell view of chromosome shape across many species and individual cells. That said, traditional heavy metal staining provides little internal chromosomal contrast, limiting insight into how individual DNA contours are organized in 3D. DNA-specific electron microscopy stains [111] could help bridge this gap, as could integrating high-resolution 3D light or electron microscopy with sequence-level information — through telomere-specific [112–115] or TAD-specific [21, 32, 33] FISH probes. These future efforts may also reconcile discrepancies between FIB-SEM and Hi-C derived structural models [61]. More broadly, our work shifts attention away from whether dinoflagellate chromosomes are helical or composed of stacked flat discs — they are predominantly neither, as most lack surface ridges entirely. Attention should instead turn to the questions that remain: What precisely are toroids made of? How many chromosomes actually are there? And what drives their vast structural heterogeneity?

## METHODS

### Dinoflagellate cell culture

*S. microadriaticum* cultures (strain # CCMP2467) were obtained from the National Center for Marine Algae and Microbiota (NCMA) at the Bigelow Laboratory for Ocean Science. *B. minutum* cultures were a gift from Dr. Annika Guse. *F. kawagutii* cultures were a gift from Dr. David Morse. *C. cohnii* cultures (strain # CCMP316) were obtained from NCMA.

Photosynthetic dinoflagellates were cultured in Daigo’s IMK medium (Fujifilm) dissolved in artificial seawater (37 g of Tropic Marin Classic sea salt mix per 1 L ddH_2_O) at 25°C, 50% humidity, with 8 am-8 pm 25 *µ*mol/(s m^2^) light exposure. A few milliliters of culture were split into 2 L of fresh IMK medium once a month.

Heterotrophic dinoflagellates were cultured in a medium consisting of 4 g/L yeast extract and 9 g/L glucose in artificial seawater. Cultures were grown at 20°C in the dark and split 1:10 once every 7-11 days.

### FIB-SEM sample preparation

Live cells were concentrated by centrifugation and cry-ofixed using high-pressure freezing (−196°C and 2150 bar, Leica EM Ice), followed by freeze substitution (Leica EM ASF2) using a mixture of 1% osmium tetroxide and 1% uranyl acetate in anhydrous acetone. Samples were gradually warmed from −140°C to −90°C over 12 h, and then held at −90°C for 72 h. Specimens were then embedded in Epoxy resin (Electron Microscopy Sciences), cured at 60°C, and trimmed using a block trimmer (Leica EM Rapid Specimen Trimming Device) such that cells were embedded just below the surface. Block faces were subsequently polished using a diamond knife (Diatome Histology) to obtain an ultra-smooth surface revealing subsurface cells. Finally, trimmed blocks were affixed to SEM stubs with silver paste and sputter-coated (Leica ACE600) with 4 nm of platinum to ensure conductivity.

### FIB-SEM imaging

High-resolution FIB-SEM nanotomography acquisitions of dinoflagellate cells were conducted at Fibics Incorporated (Ottawa) using a Zeiss Crossbeam 550 FE-SEM (*F. kawagutii*) and a Zeiss Crossbeam 350 (*S. microadriaticum, B. minutum*, and *C. cohnii*). All Zeiss datasets were acquired using the Atlas 3D (A3D) Nanotomography module of the Zeiss Atlas 5^TM^ software. For the A3D run preparation, an initial protective layer of platinum (~ 1 *µ*m) was deposited on top of the volume to be imaged before 3D tracking and SEM autotune patterning. The marks were highlighted with a thin carbon deposit followed by a second protective layer of carbon (~ 1 *µ*m). A cross-section face for A3D imaging was exposed by creating a two-stage stepped trench using a 30 nA beam current for the coarse trench followed by 3 nA beam current to remove the remaining material up to the starting cross-section face of the volume to be imaged. Automated FIB milling was carried out with the Ga^+^ FIB at an acceleration voltage of 30 kV and a 1.5 nA beam current. Automated SEM image acquisition was done with both the InLens and SE2 detectors at an acceleration voltage of 2 kV and a probe current of 500 pA. During all A3D acquisitions, an hourly automatic correction of focus and astigmatism was performed.

*F. kawagutii* and *B. minutum* cells were acquired using the “Conventional” imaging and acquisition mode with the option “pause milling while imaging” selected. We used a 4 nm target slice thickness, a pixel size of 4 nm, and dwell times of 2 *µ*s for *F. kawagutii* and 3 *µ*s for *B. minutum*. The resulting images were aligned within Atlas 5 and exported as TIFF image stacks. Datasets for *C. cohnii* and *S. microadriaticum* were acquired using the “Thin & Fast” imaging and acquisition mode with the option “continuous milling and imaging” selected. We used a target slice thickness of 1 nm, a 4 nm pixel size, and a 0.5 *µ*s dwell time for both species. These datasets were aligned within Atlas 5 and exported as TIFF image stacks using the Intelligent Integer Integration algorithm with a binning of 4. Imaging continued until at least three separate nuclei were imaged for each species, often including whole or significant fractions of cells.

Due to instrument availability, one *C. cohnii* cell was imaged at the Microscopy and Material Characterization Center at Polytechnique Montréal on a Thermo Fisher Helios 5 Hydra CX FIB-SEM. Sample preparation involved depositing a 2 *µ*m platinum protective pad, cutting a step trench directly in front of the pad using oxygen plasma (30 kV, 60 nA), and polishing the sample crosssection using 3.2 nA oxygen plasma. Automated FIB milling was then performed again with oxygen plasma at 30 kV but with a higher current (1.7 nA). SEM images were acquired using the Through-the-Lens Detector at 2 kV, 400 pA, and a 30 *µ*m aperture, with a slice thickness and pixel size of 4 nm and a dwell time of 4 *µ*s.

### Image processing and deep learning segmentation

FIB-SEM data originating from the literature were downloaded from either the EMPIAR or BioImage Archive databases using accession codes: EMPIAR 47483651 [40], BioImage S-BSST575 [39], EMPIAR 11399 [41], and EMPIAR 12627 [42].

All image processing and segmentation were done using Dragonfly v.2024.1 (Comet Technologies Canada Inc.). FIB-SEM image stacks were loaded in batches with the same X/Y dimensions and stitched together using Dragonfly’s 3D Image stitching tool. Slice registration was accomplished using a sum of squared differences algorithm (max step 2%, no rotation). Horizontal and vertical de-striping was accomplished using a wavelet transform using default parameters (*σ* _foreground_ = 128 pixels, *σ* _background_ = 256 pixels, Levels = 7, Wavelet = db21). 3D gaussian smoothing was then performed (kernel = 3 voxels, *σ* = 1.2), and a box crop was done enclosing the nucleus. For 5-8 slices throughout the nuclear subvolume, chromosomes were manually labelled using Dragonfly’s built-in paint tools. This “ground truth” was used to train a U-Net learning classifier (initial filter count = 512, depth = 5 layers, patch size = 64, batch size = 256, optimization algorithm = Adadelta, Loss function = CategoricalCrossentropy) that was applied to the whole image stack. A separate deep learning model was trained for each nucleus. A multi-region of interest (multi-ROI) was created for each class using connected components (6-connected) and objects outside the nucleus were discarded. A final round of manual segmentation was performed to eliminate minor segmentation errors. A custom menu option was developed that reorients each ROI within a multi-ROI, i.e. each chromosome, such that its long axis now points vertically (along z) and exports each oriented ROI as a 3D binary TIFF file (Code availability).

### Spherical Harmonics Expansion (SHE) and Principal Component Analysis (PCA)

Code implementing SHE and PCA is hosted in our GitHub repository (Code availability). This code relies on the aics-shparam Python package [51].

To maintain reconstruction accuracy, chromosomes were omitted if they exceeded a reconstruction error threshold. As done previously [51], we defined reconstruction error, *RE*, as

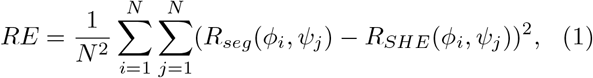

where *R* is the radius at a particular longitude (*ϕ* ∈ [0, 2 *π*]) and latitude (*ψ* ∈ [0, *π*]) coordinate for either the original segmentation, *R*_*seg*_, or the Spherical Harmonic Expansion, *R*_*SHE*_. *N* ^2^ total points were used for each mesh, corresponding to the latitude and longitude coordinates where *R*_*SHE*_ was optimized to match *R*_*seg*_.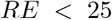, which guarantees an average error less than 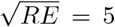 voxel heights across all mesh points. Chromosomes containing large holes or whose centroid is located outside its volume were omitted. *F. kawagutii* chromosome networks were omitted as chromosomes are interconnected and not individually segmentable.

The parameter *l*_*max*_ determines the number of spherical harmonic functions, 2(*l*_*max*_ + 1)^2^, used in the expansion [51]. We found *l*_*max*_ = 40 generally yielded chromosomes with a low reconstruction error (Fig. S5 a).

Given our choice of *l*_*max*_ = 40 and *RE* < 25, 1810 of 2138 total chromosomes (85%) were reconstructed successfully and included in our analysis.

### Voronoi simulations

Voronoi simulations were performed using Dragonfly’s built-in Python console. All segmentations — including chromosomes, nucleoli, and the nuclear membrane — were downsampled from 4 nm to 40 nm to reduce the simulation run-time. Chromosome centroids were then iteratively dilated using a previously developed kernel [116] until the nuclear volume was completely filled. A detailed procedure is available in our GitHub repository (Code availability).

### Surface ridge angle extraction

Vertically-oriented 3D chromosome TIFF files were split into front half and back half volumes and projected inwards onto the bisecting plane using a custom MATLAB script. Front half and back half projections were saved as separate images and the protractor tool in Fiji was used to manually extract surface ridge angles relative to the long axis. Manually extracted angles were verified using a custom MATLAB script, which generated overlay galleries on front- and back-half chromosome projections; any angles that failed visual inspection were re-measured.

### Skeletonization

Prior to skeletonization, Dragonfly’s “Close” function (kernel = 7 voxels) and “Fill inner areas” were applied to eliminate artifactual porosities within the *F. kawagutii* chromosome network ROI. The 3D binary ROI was exported as a TIFF file, then converted to an MRC file using tif2mrc from IMOD [117]. We used Voxel Cores’ [58] vol2mesh command setting onlyBndryVts=false to produce an OFF boundary mesh file including faces. Voxel Cores was then used to generate a preliminary medial axis skeleton (PLY file) and a corresponding radial distance file (R file) using the vol2ma command with a pruning threshold of *λ* = 0.006. The skeleton was further pruned using Erosion Thickness [57] with a maximum “ET on MA” (erosion thickness on medial-axis) threshold, and a threshold of 25 for “ET on MC” (erosion thickness of medial-curve). Erosion Thickness outputs the final pruned skeleton (PLY file) along with the maximally inscribed sphere radius, *r*, at each skeleton point.

The *F. kawagutii* chromosome skeletons were subdivided into parts using a threshold of *r*_*thresh*_ = 130 nm. Parts of the skeleton with an inscribed sphere radius *r >* 130 nm were designated as thick tubes while parts of the skeleton with a *r* < 130 nm were designated thin bridges. These skeleton parts were subdivided further at junction nodes with connectivity degree ≥ 3. Each skeleton part was visually inspected using the napari Python package [118] (Code availability) for quality and its contour length was measured. Contour length distributions were visualized using a “Superplot” [119].

### Voxel intensity analysis

Using Dragonfly, 3D segmentations were used as masks to extract voxel intensities, *I*, within each object type, *i*, yielding normalized intensity histograms, *g*_*i*_(*I*), satisfying 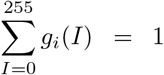. The summation range *I* ∈ *{*0, 1,…, 255*}* corresponds to the possible intensity values for an 8-bit image. We defined the Percent Intensity

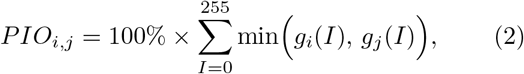

where *i* corresponds to all of: rods, crescents, and toroids, while *j* corresponds to one of: chromosomes, nucleolus, or nucleoplasm. *PIO*_*i,j*_ was computed separately for each cell, then averaged across cells while holding *i* and *j* fixed to obtain 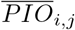.

## DATA AVAILABILITY

Raw FIB-SEM images are available in the Electron Microscopy Public Image Archive (EMPIAR) using the accession code: EMPIAR-13824. The aligned FIB-SEM images and segmentations can be interactively explored via the ATLAS browser-based viewer—currently available as a prototype—at: https://petapixelproject.com/urls/philipp2026.

Data and analysis files have been uploaded to Zenodo: https://zenodo.org/records/21478140. These include: a spreadsheet of volume measurements; 3D binary TIFF files of vertically oriented chromosomes; 3D binary TIFF files of rods, crescents, and toroids; image galleries of extracted surface ridge angles (SI Data 1-21); 3D videos of *F. kawagutii* chromosome network segmentations (SI Videos 1-3); and *F. kawagutii* skeletonization PLY files.

## CODE AVAILABILITY

Code for reproducing the analyses presented here are hosted on our GitHub repository: https://github.com/lucasphilipp1/3D-Electron-Microscopy-of-Dinoflagellate-Chromosomes-Across-Many-Species.

## ACKNOWLEDGMENTS

We would like to thank Dr. David Morse (Université de Montréal) and Dr. Annika Guse (Ludwig-Maximilians-Universität München) for providing dinoflagellate starter cultures; Dr. Kelly Sears (McGill) for maintaining the Facility for Electron Microscopy Research (FEMR) core facility; Dr. Etienne Bousser (Polytechnique), Dr. Weawkamol Leelapornpisit (FEMR, McGill), Dr. Stéphanie Bessette (FEMR, McGill), and Dr. Angel Valdez (FEMR, McGill) for assistance operating the FIB-SEMs, and for training; Benjamin Rudski and Dr. Joseph Deering for their expertise using Dragonfly; Dr. Marc McKee for equipment access and useful discussions; and Dr. Abigail Gerhold for access to a Windows 11, 125 GB RAM, GPU (Intel UHD Graphics 770 & NVIDIA RTX 2000 Ada Generation) workstation. We also thank Comet Technologies for providing a free-of-charge academic license for Dragonfly.

This work was funded by the New Frontiers in Research Fund (NFRFE-2019-00189 to S.C.W.) and by a CRBS Blue Sky Seed Funding Award from the Centre de Recherche en Biologie Structurale, funded by Fonds de Recherche du Québec (Health Sector) Grant (#288558 to S.C.W. and N.R.). L.P. was supported by a Canadian Graduate Scholarship – Master’s and a Canadian Graduate Scholarship – Doctoral from the National Science and Engineering Research Council of Canada. E.I. is a recipient of a Vadasz Doctoral Fellowship. N.R. is a William Dawson Scholar at McGill University. This research was undertaken, in part, thanks to funding from the Canada Research Chairs Program (to S.C.W.).

## AUTHOR CONTRIBUTIONS

L.P. and S.C.W. designed the research and wrote the paper; L.P., S.C.W., and N.R. obtained funding; N.R. provided experimental and image analysis training; L.P. and E.I. prepared algae samples for imaging; D.S., E.I., and L.P. performed FIB-SEM; J.F. developed the browser-based data viewer; L.P. performed all other aspects of the research.

