## Supplementary Information for "3D Electron Microscopy Reveals Diverse Chromosome Morphologies Across Dinoflagellate Species"

### SUPPLEMENTARY INFORMATION (SI)

Supplementary Information accompanying this article including SI Figures S1-S10 and SI Table 1 can be found below. See Data Availability in main text for SI Data 1-21 and SI Videos 1-3.

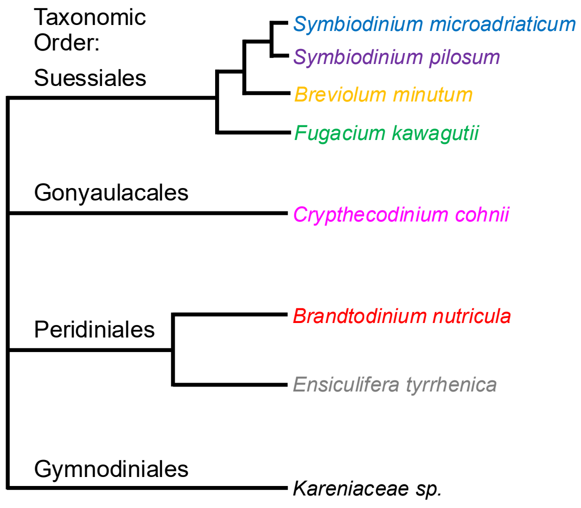


FIG S1. Qualitative relatedness of dinoflagellate species analyzed in this study (see Table I).

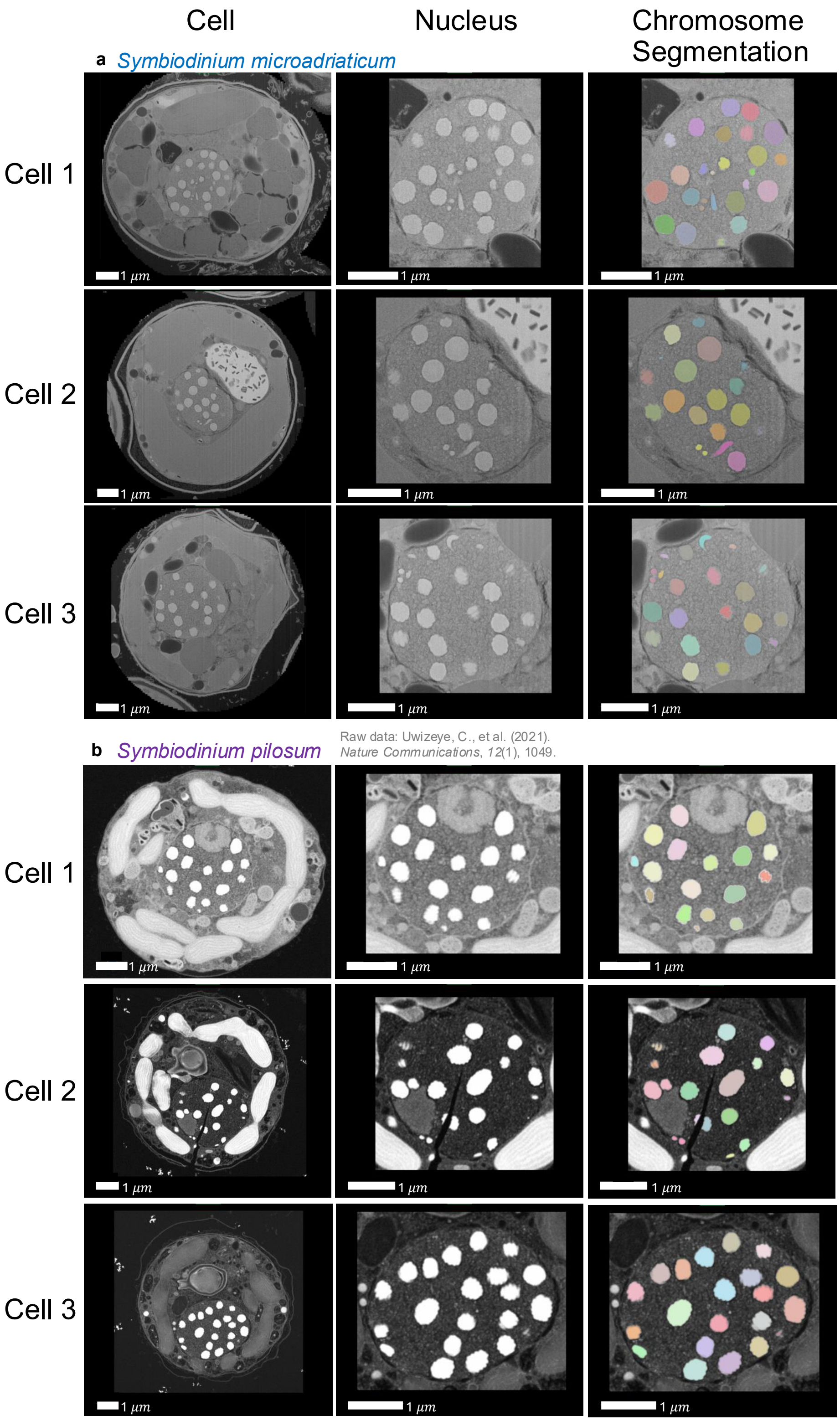


FIG S2. FIB-SEM images of cells, nuclei, and nuclei with segmented chromosomes. Each row is an individual cell. Sources of raw images where not original are indicated.

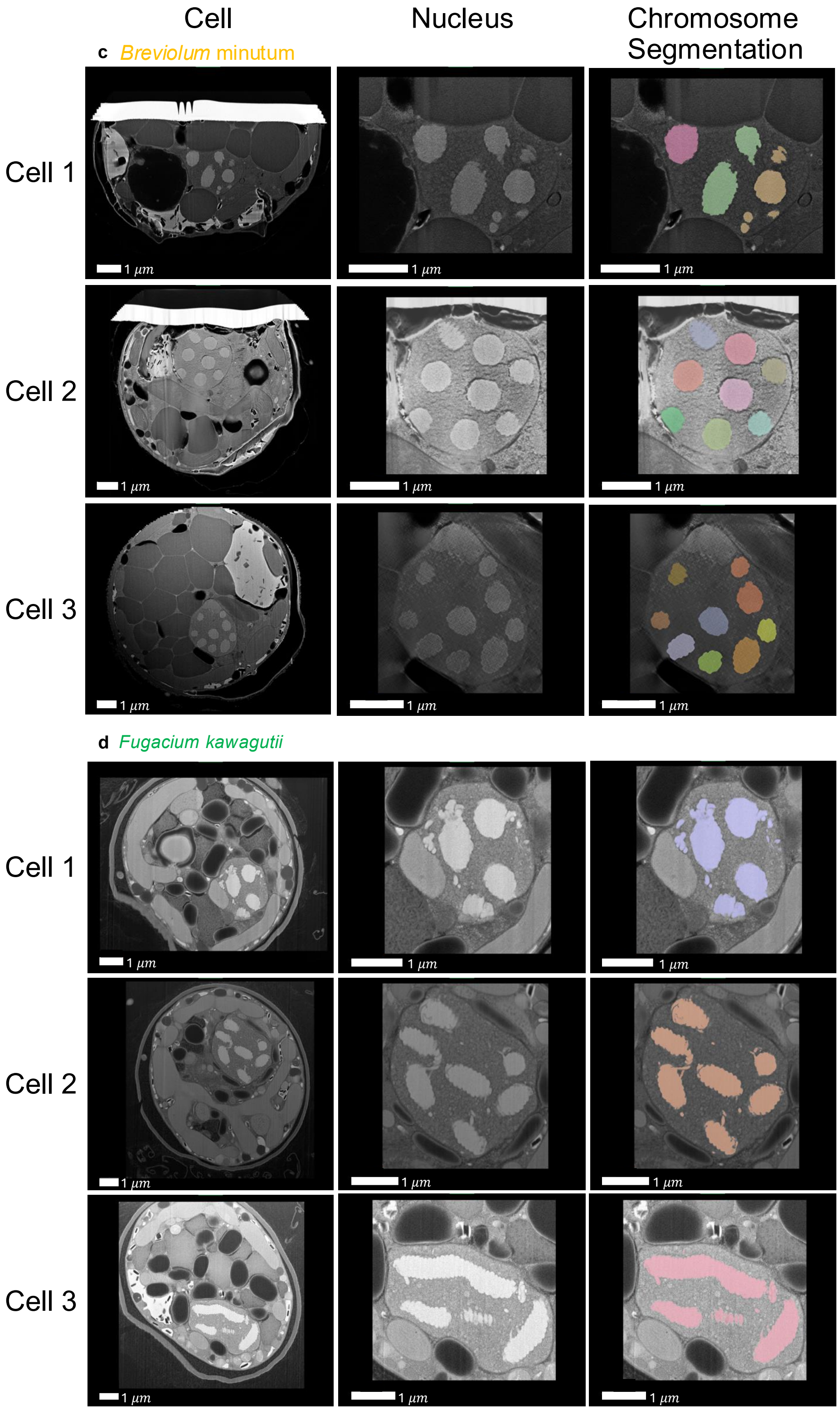


FIG S2. FIB-SEM images of cells, nuclei, and nuclei with segmented chromosomes. Each row is an individual cell. Sources of raw images where not original are indicated.

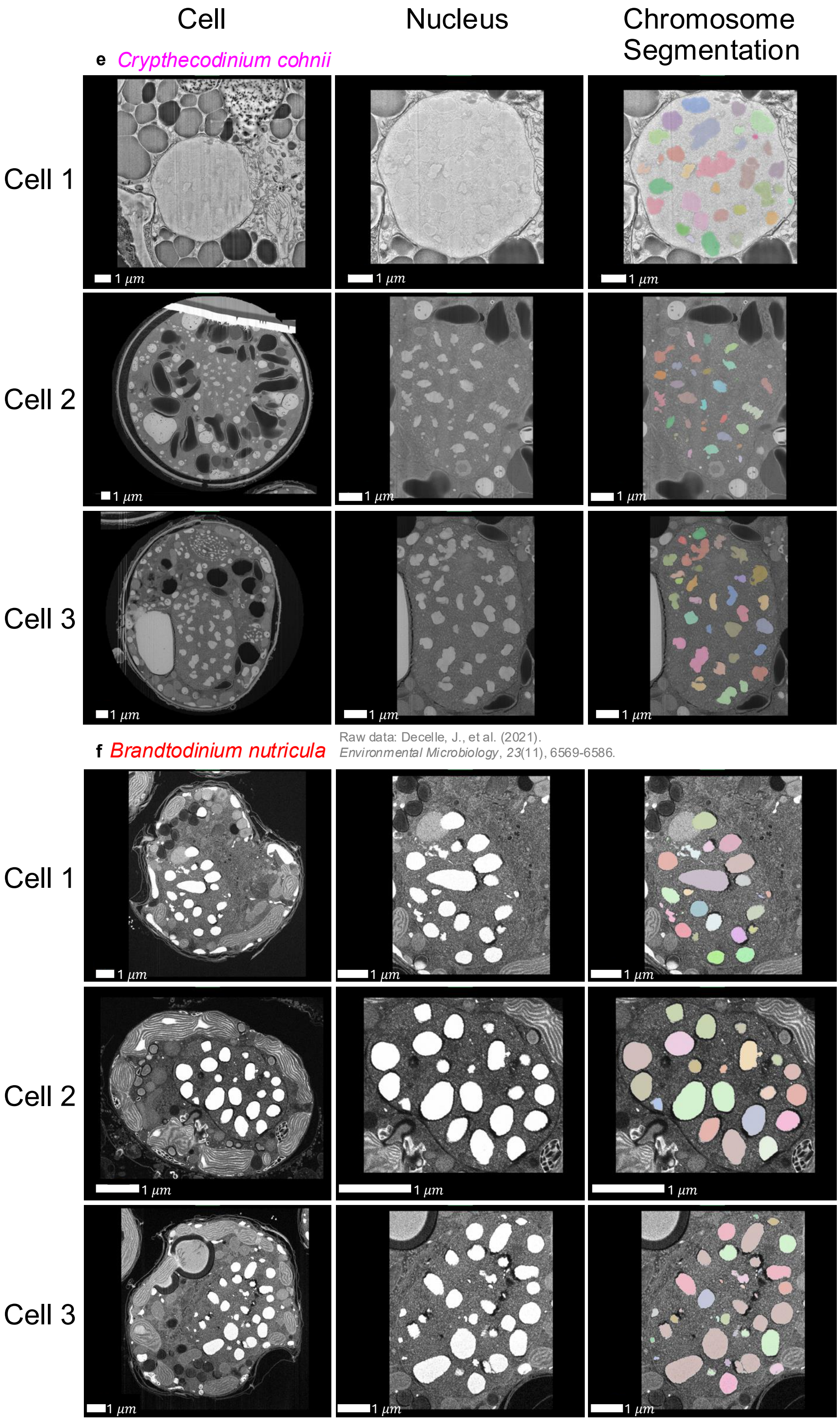


FIG S2. FIB-SEM images of cells, nuclei, and nuclei with segmented chromosomes. Each row is an individual cell. Sources of raw images where not original are indicated.

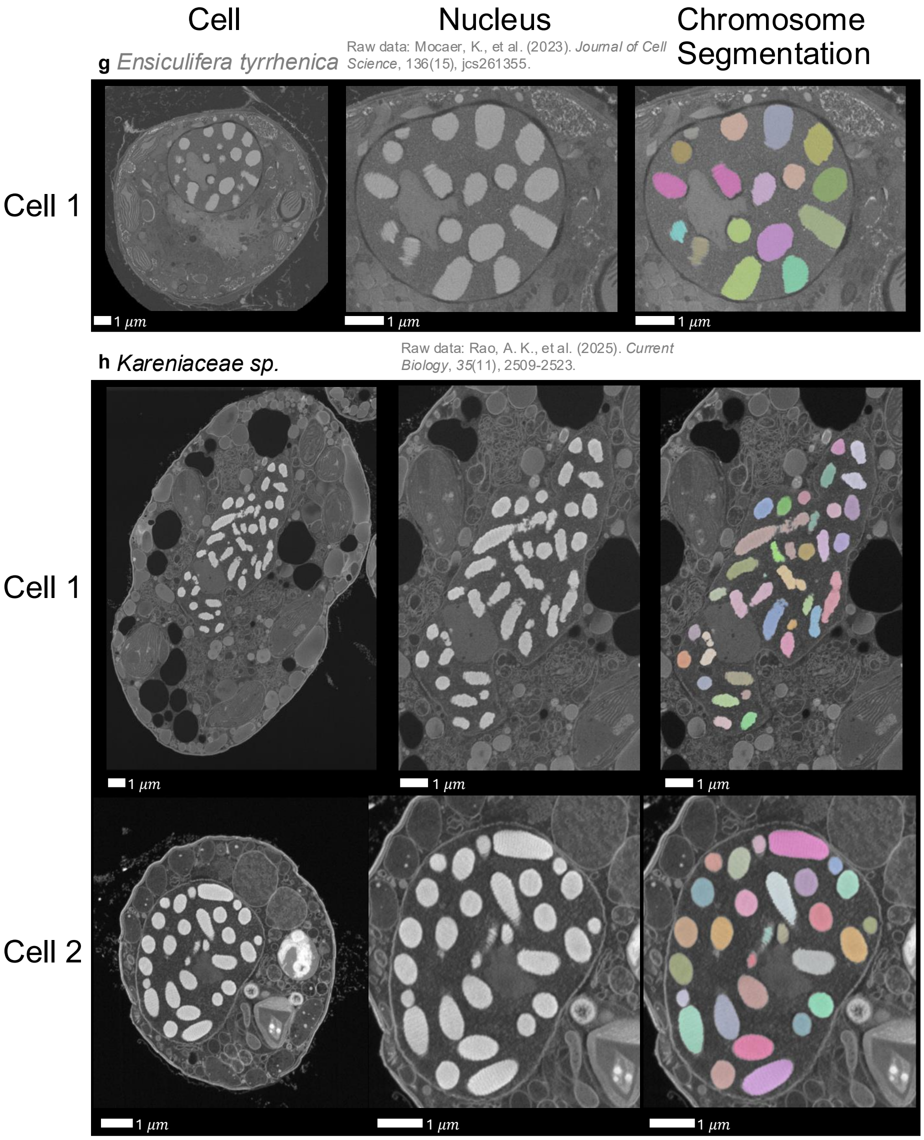


FIG S2. FIB-SEM images of cells, nuclei, and nuclei with segmented chromosomes. Each row is an individual cell. Sources of raw images where not original are indicated.

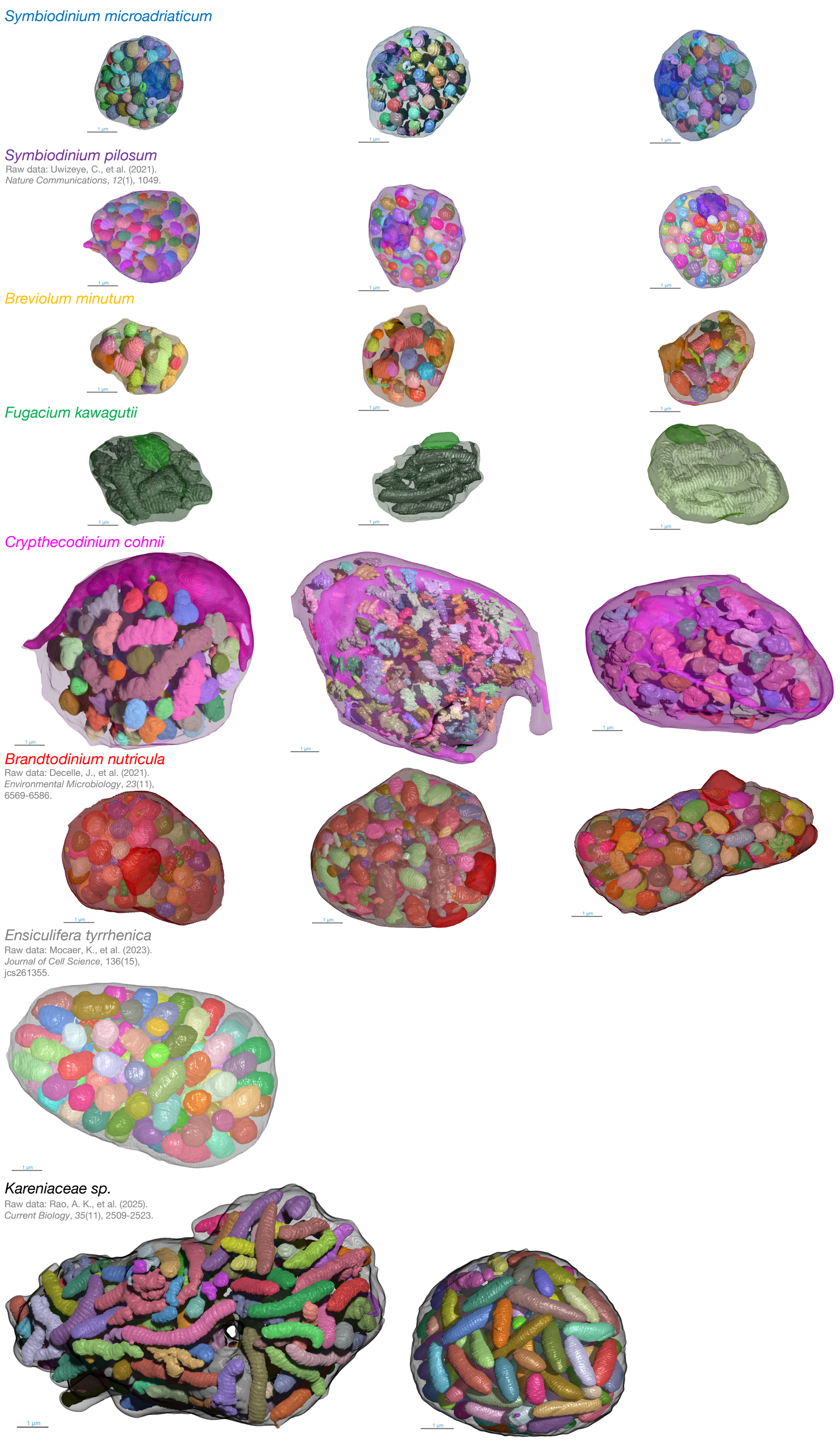


FIG S3. To-scale 3D reconstructions of all dinoflagellate nuclei analyzed in this study (see Table I). Chromosomes, nucleoli, and the nuclear membrane are shown. Each object is assigned a random color for better distinguishability. Sources of raw images where not original are indicated. Note the hole through the nucleus in *Kareniaceae sp.* Cell 1.

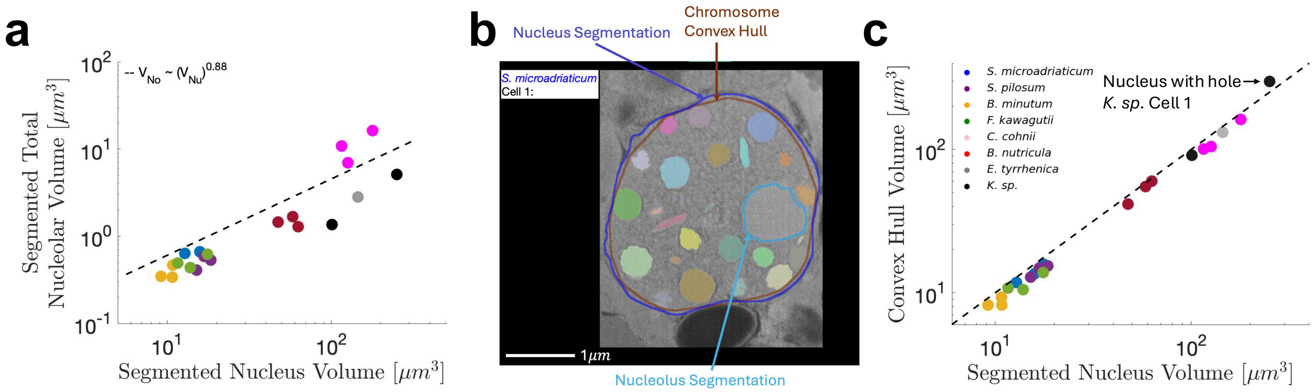


FIG S4. a) Segmented total nucleolar volume*, $V_{No}$, increases with segmented nuclear volume, $V_{Nu}$, non-linearly. Dashed line represents the best power-law fit. b) Visualization of the segmented nucleus and nucleolus, as well as the convex hull containing all chromosomes. c) The volume of the convex hull containing all chromosomes is an accurate approximation ($R^2 = 0.985$) of the segmented nuclear volume. If chromosomes are already segmented, computing the convex hull is straightforward while segmenting the nucleus requires training another deep learning model [1]. The convex hull measurement is typically an underestimate, except for nucleus from *K. sp.* Cell 1, which has a hole through it (see Fig. S3). *Some nuclei have > 1 nucleolus, in which case the volumes of individual nucleoli were summed.

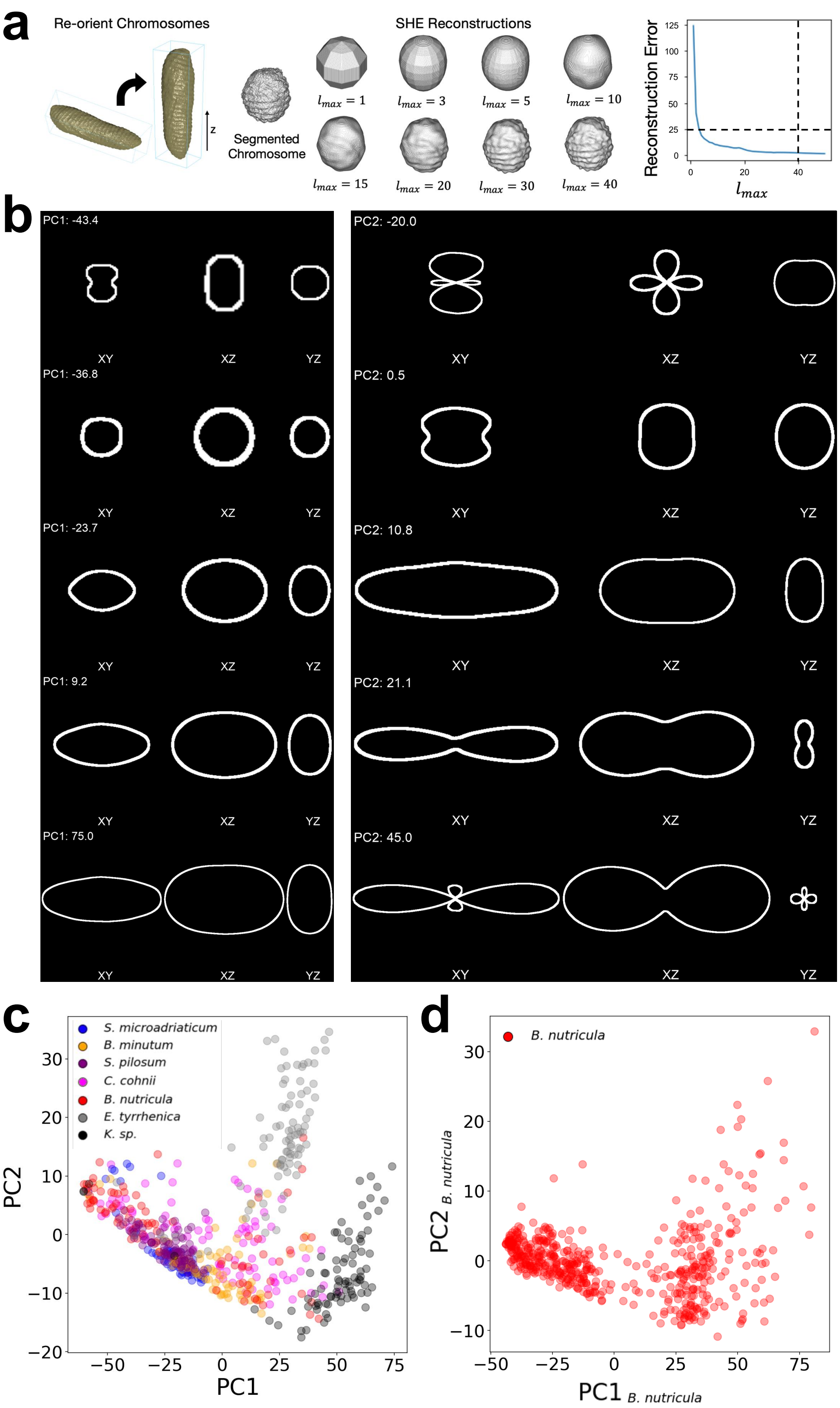


FIG S5. a) Prior to SHE, chromosomes were re-oriented so their long axis points vertically. That way the SHE depends only on chromosome shape, not orientation. The number of spherical harmonics in the expansion increases monotonically with $l_{max}$. Higher $l_{max}$ typically results in lower reconstruction error. Dashed lines indicate the thresholds selected for $l_{max}$ and maximum reconstruction error tolerated for SHE. b) Cross-sections of shapes generated by sampling the principal component axes of shape space. Left: varying $PC1$, setting $PC2 = 0$. Right: setting $PC1 = 0$, varying $PC2$. c) PCA using an equal number of chromosomes for each species recapitulates Fig. 3 e. d) PCA using only *B. nutricula* chromosomes is highly similar to the PCA using all chromosomes (Fig. 3 e). There are two chromosome populations, one with small volumes and one with large volumes.

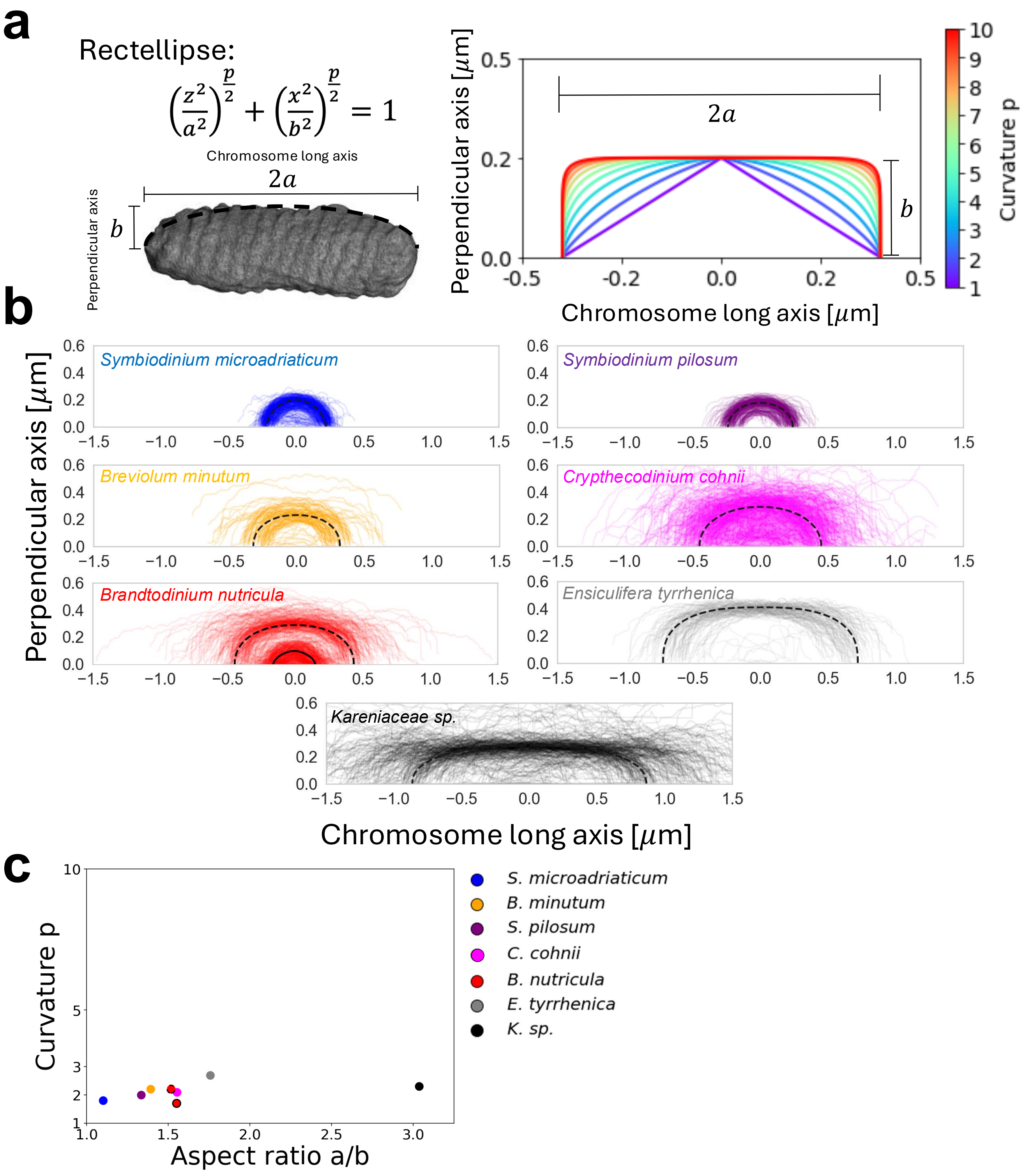


FIG S6. a) The rectellipse equation parametrizes chromosomes of different length, width, and curvature profiles. $z$ is the horizontal (long) axis, $x$ is the perpendicular axis, $2a$ is the chromosome length, $2b$ is the chromosome width, and $p$ is the curvature. b) Cross-sectional outlines of chromosomes with their long axis aligned horizontally. Each curve is a chromosome. Dashed arcs are rectellipse approximations. For *B. nutricula*, there are two chromosome populations, one small (solid line) and one large (dashed line). c) Curvature and aspect ratio for each species taken from rectellipse approximations. Dinoflagellate chromosome curvature is largely independent of aspect ratio. *B. nutricula* dot outlines correspond to arcs in b.

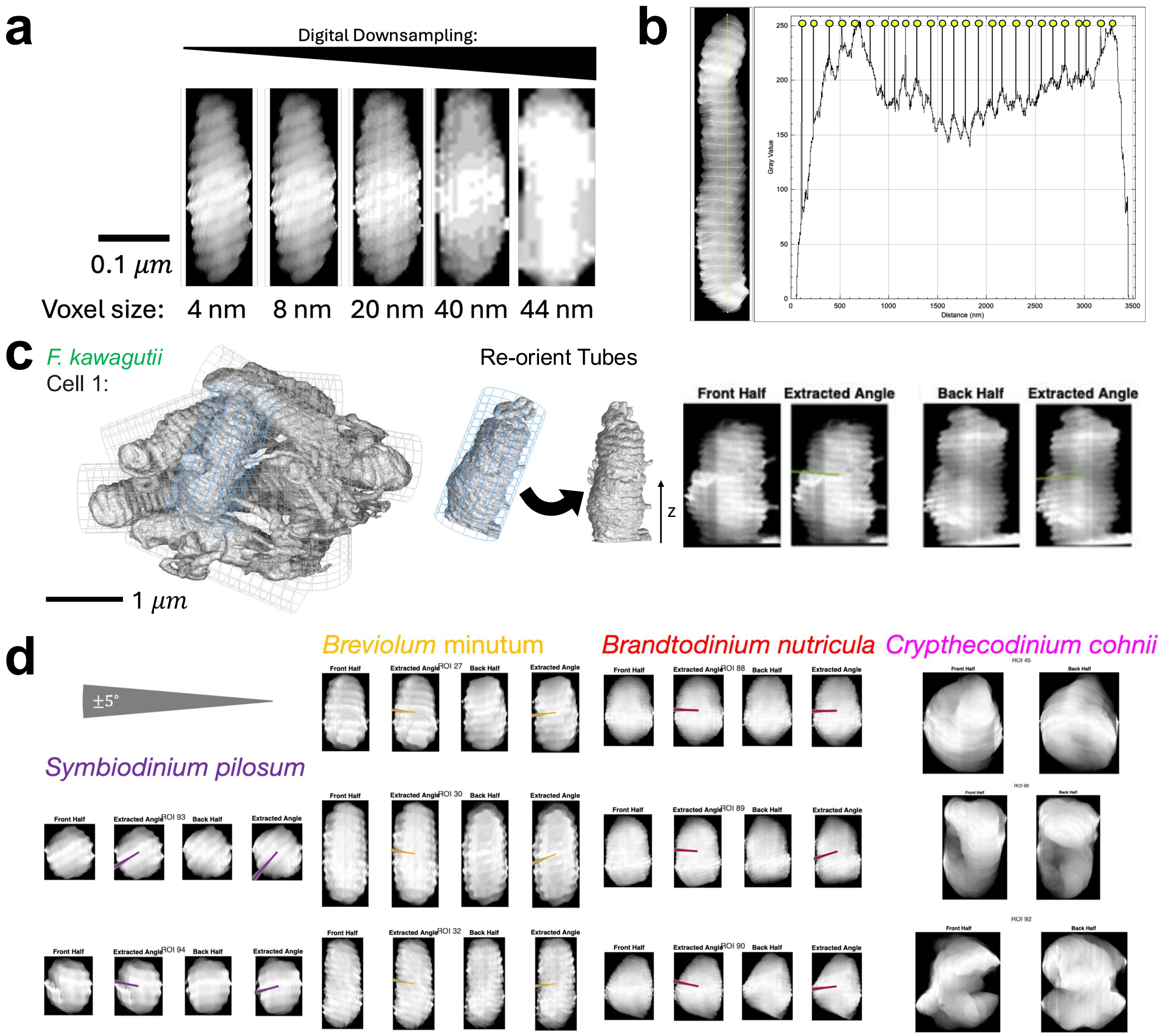


FIG S7. a) Detection of surface ridges requires sufficiently high resolution, which was not attained in previous studies [2, 3]. b) Ridges were manually annotated using the draw tool in Fiji [4]. Their locations (yellow circles) were then extracted using a line-scan by thresholding the intensity at the maximum value. c) For species with isolated chromosomes, a bounding box surrounding each chromosome was used to re-orient it so that its long axis pointed vertically (see Fig. S5 a). For *F. kawagutii* chromosome networks, a cylindrical cropping region/bounding box was placed around each tube and used to re-orient that genomic segment so its long axis points vertically. Then after projecting the genomic segment onto its middle plane, front half and back half surface ridge angles were extracted. d) Verification of ridge angle measurements was done via visual inspection. A $\pm 5^\circ$ angle tolerance is shown, accounting for labelling uncertainties. Example chromosomes from four randomly chosen species are shown. Surface ridges in *C. cohnii* chromosomes are less common. For complete galleries showing all chromosomes from all cells and species, see SI Data.

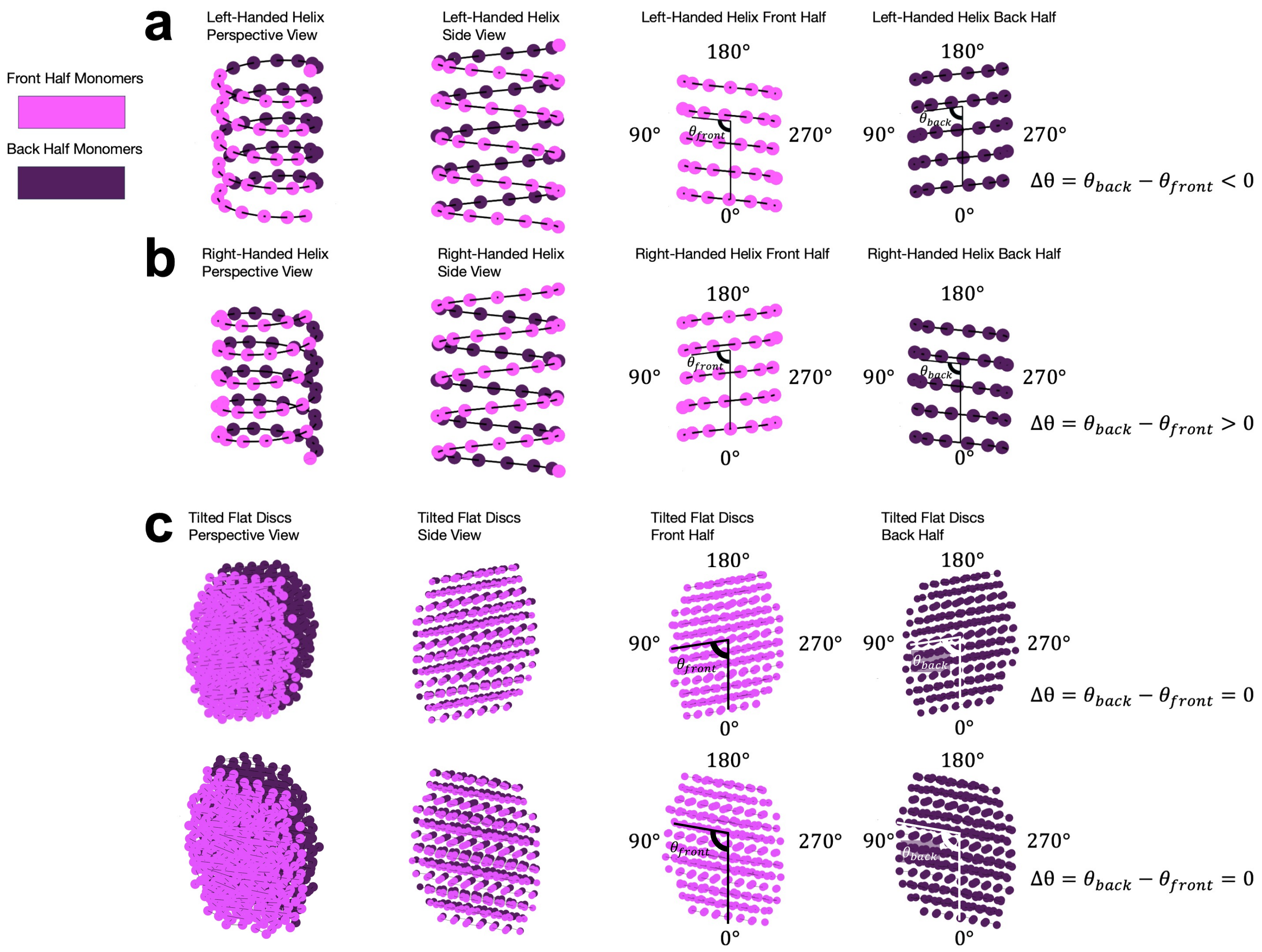


FIG S8. $\Delta\theta = \theta_{back} - \theta_{front}$ distinguishes a) left-handed helices, b) right-handed helices, and c) stacks of flat discs regardless of tilt. For each category, monomers are colored by their location: front half (pink) or back half (purple). Polymer bonds are shown in black. Perspective (left) and side (center left) views are shown, with extracted angle from the front half (center right) or back half (right) projections indicated.

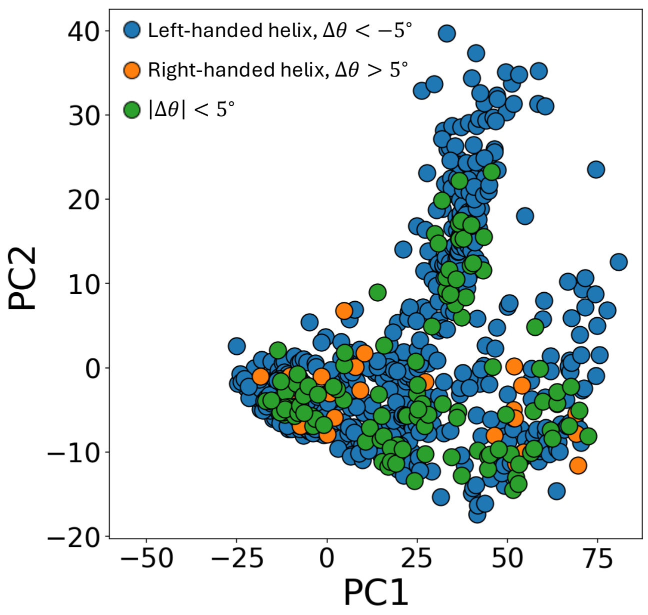


FIG S9. There is significant overlap in PCA shape space between chromosomes with left-handed helical ($\Delta\theta < -5^{\circ}$) and flat disc ($|\Delta\theta| < 5^{\circ}$) organization. Chromosomes with right-handed helical organization have small aspect ratios (lower/more negative $PC2$ values).

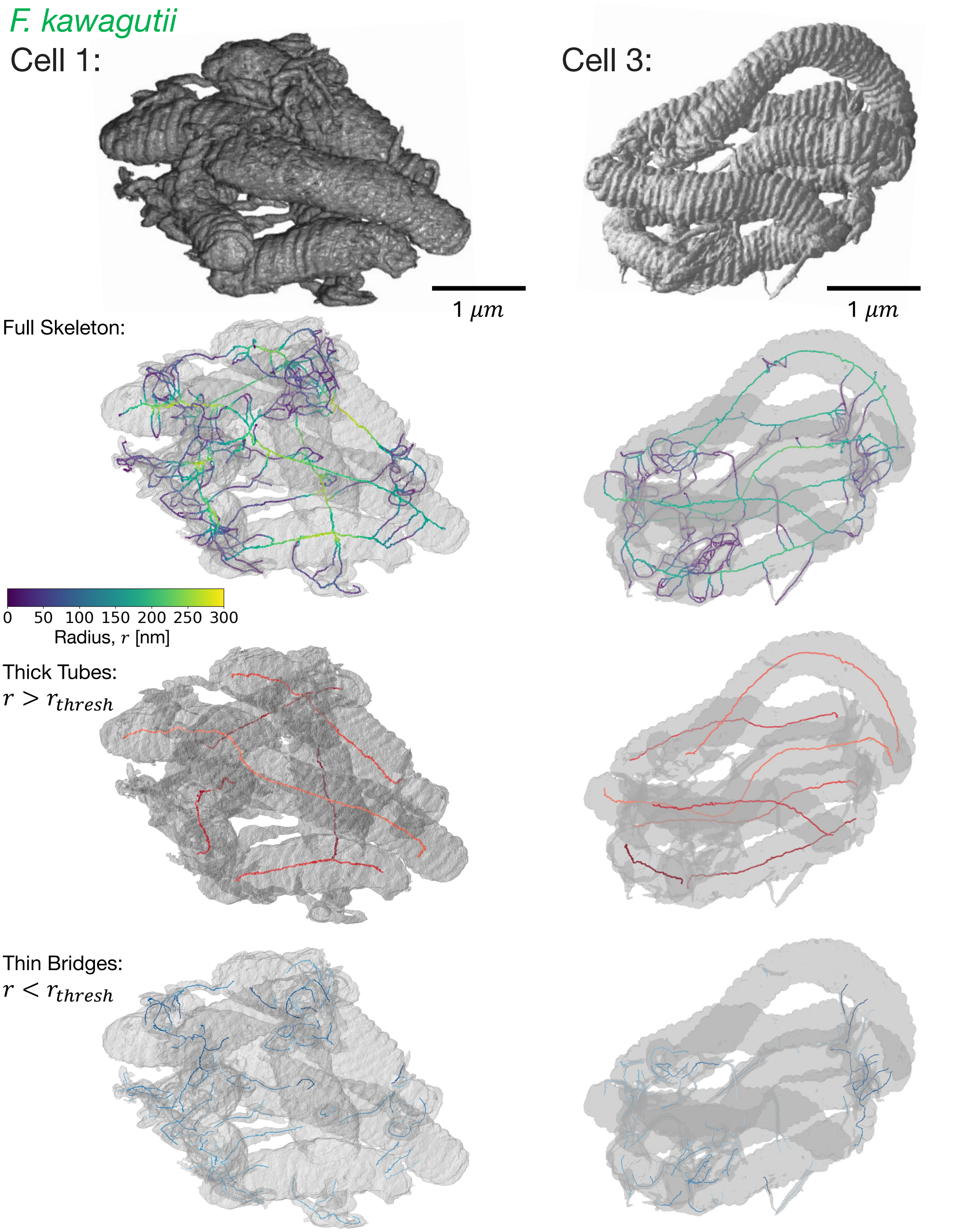


FIG S10. Skeletonization of *F. kawagutii* Cells 1 and 3. Colorbar indicates the radius, $r$, of the largest inscribed sphere centred at points along the skeleton. Visualization of skeleton parts separated as thick tubes ($r > r_{\text{thresh}}$) or thin bridges ($r < r_{\text{thresh}}$), where $r_{\text{thresh}} = 130$ nm. Each thick tube and thin bridge is assigned a random color shade for better distinguishability.

TABLE S1. Summary of observed genomic and morphological features across dinoflagellate species

| Order | Species | Genomic features[a] | | | | Morphological features | | |
|---|---|---|---|---|---|---|---|---|
| | | Length | DVNPs [5–7] | HLPs [8] | Tandem repeats | Technique | Sample prep. | Exceptions[b] |
| Suessiales | ***S. micro-adriticum***[c] | 1.10 Gb | Yes | | 50% of genes in $\geq 9\times$ arrays [9] | FIB-SEM | HPF/FS[d] | No exceptions. |
| | ***S. pilosum*** | 1.99 Gb | Yes | | | FIB-SEM[10] | HPF/FS | ‖[e] |
| | *Symbiodinium* clade A | | Yes | HLP-I | | TEM[11] | CF[f] | No arches. |
| | ***B. minutum*** | 1.50 Gb | Yes | HLP-I | 4.6% of genome [12] | FIB-SEM | HPF/FS | No exceptions. |
| | *Symbiodinium* clade C | | Yes | HLP-I | | TEM[13] | CF | No arches. |
| | ***F. kawagutii*** | 1.18 Gb | Yes | | 2% of genes in $\geq 4\times$ arrays [14] | FIB-SEM | HPF/FS | One contiguous chromosome network. |
| | *A. granifera* | | | | | TEM[15] | CF | |
| | *B. andersenii* | | | | | TEM[16] | CF | No surface ridges. No arches. |
| Gonyaulacales | ***C. cohnii*** | 25 Gb | Yes | HLP-I | | FIB-SEM | HPF/FS | No exceptions. |
| | | | | | | TEM[17] | CF | Some uncoiling of surface ridges. |
| | | | | | | TEM[18] | CF | No exceptions. |
| | *G. polyedra* | | | | | TEM[19] | HPF/FS | ‖ |
| | *A. tamarense* | | Yes | | | TEM[20] | CF | ‖ |
| Peridiniales | ***B. nutricula*** | | Yes | | | FIB-SEM[21] | HPF/FS | ‖ |
| | ***E. tyrrhenica*** | | | | | FIB-SEM[22] | HPF/FS | ‖ |
| | *C. arcachonensis* | | | | | TEM[23] | CF | No arches. |
| | *P. cinctum* | | | | | TEM[24] | CF | Uncoiling surface ridges. |
| | *H. claromecoensis sp.* | | | | | TEM[25] | CF | No exceptions. |
| | *P. balticum* | | | | | TEM[26] | CF | Two nuclei: dinokaryon & eukaryotic |
| | *H. triquetra* | | | | | TEM[20] | CF | No exceptions. |
| | *D. kwazulunatalensis* | | | | | TEM[27] | CF | Two nuclei: dinokaryon & eukaryotic |
| | *D. capensis* | | | | | TEM[27] | CF | Two nuclei: dinokaryon & eukaryotic |
| Prorocentrales | *P. micans* | | | | | TEM[28] | HPF/FS & CF | No exceptions. |
| | | | | | | TEM[20] | CF | ‖ |
| | | | | | | TEM[29] | CF | ‖ |
| | | | | | | TEM[30] | CF | ‖ |
| | | | | | | TEM[31] | CF | ‖ |
| | | | | | | TEM[32] | HPF/FS & CF | ‖ |
| | *P. minimum* | | Yes | HLP-I | | TEM[33] | CF | No arches. |
| | *P. cordatum* | | | | | FIB-SEM[2] | CF | No surface ridges. |

*Continued on next page...*

*Continued from previous page...*

| Order | Species | Genomic features[a] | | | | Morphological features | | |
|---|---|---|---|---|---|---|---|---|
| | | Length | DVNPs [5–7] | HLPs [8] | Tandem repeats | Technique | Sample prep. | Exceptions[b] |
| | ***K. sp.*** | | Yes | | | FIB-SEM[34] | HPF/FS | No exceptions. |
| | *K. brevis* | | Yes | HLP-I | | TEM[20] | CF | ‖ |
| | *K. veneficum* | | | HLP-I | | TEM[35] | CF | ‖ |
| | *A. carterae* | | Yes | HLP-II | | TEM[20] | CF | ‖ |
| Gymnodiniales | *T. magnum* | | | | | TEM[36] | CF | No surface ridges.<br>No arches.<br>No bands. |
| | *T. testudo* | | | | | TEM[37] | CF | ‖ |
| | *T. maedaense* | | | | | TEM[37] | CF | ‖ |
| | *T. corrugatum* | | | | | TEM[37] | CF | ‖ |
| | *P. jejuensis* | | | | | TEM[38] | CF | ‖ |
| Oxyrrhinales | *O. marina* | | Yes | | | FIB-SEM & TEM [3] | HPF/FS & CF | No surface ridges.<br>No arches.<br>No bands. |
| | | | | | | TEM [39] | CF | ‖ |
| Syndiniales[g] | *A. ceratii* | 0.13 Gb [40] | Yes | | | TEM [41] | CF | Not condensed.<br>Not discrete.<br>No surface ridges.<br>No arches. |
| | *S. sp.* | | | | | TEM [42] | CF | Sometimes condensed.<br>No surface ridges.<br>No arches.<br>No bands. |
| | *S. sp.* | | | | | TEM [43] | CF | Not discrete.<br>No surface ridges.<br>No arches.<br>No bands. |
| | *H. sp.* | | Yes | | | TEM [44] | CF | No surface ridges.<br>No arches.<br>No bands. |
| Noctilucales[g] | *N. scintillans* | | Yes | HLP-II | | TEM [45] | CF | As trophont:[h]<br>Not condensed.<br>Not discrete.<br>No surface ridges.<br>No arches.<br>No bands.<br>As spore:[i]<br>No exceptions. |
| Blastodiniales[g] | *B. sp.* | | | | | TEM [46] | CF | As trophont:<br>Not condensed.<br>Not discrete.<br>No surface ridges.<br>No arches.<br>No bands.<br>Sometimes as spore:<br>No surface ridges.<br>No arches.<br>No bands. |

[a] Blanks indicate data unavailable.
[b] Unless otherwise stated, chromosomes are “dinokaryonic”, i.e.: condensed, discrete, and cylindrical, with surface ridges, alternating bands of low/high electron density, and arches in oblique cross-sections.
[c] Species in bold were either imaged or analyzed in this study.
[d] HPF/FS (High-Pressure Freezing/Freeze Substitution): Cryogenic sample preparation method where water is first rapidly vitrified to prevent ice crystal formation. Amorphous ice is then substituted at ultra-low temperatures with organic solvents containing fixatives and heavy-metal stains. After fixation, the waterless sample is slowly warmed to room temperature.
[e] Same as above.
[f] CF (Conventional Fixation): Fixation, heavy-metal staining, and dehydration performed at room temperature.
[g] Obligate parasites.
[h] Trophont: parasitic life stage, feeding inside the host.
[i] Spore: parasitic life stage, flagellated and swimmming outside the host.

[23] Zhaohe Luo, Kenneth Neil Mertens, Elizabeth Nézan, Li Gu, Vera Pospelova, Hikmah Thoha, and Haifeng Gu. Morphology, ultrastructure and molecular phy-

logeny of cyst-producing *Caladoa arcachonensis* gen. et sp. nov.(peridiniales, dinophyceae) from france and indonesia. *European Journal of Phycology*, 54(2):235–248, 2019.
